# The long non-coding RNA *MaTAR20* promotes mammary tumor growth by regulating angiogenesis pathways

**DOI:** 10.1101/2021.03.30.437774

**Authors:** Sarah D. Diermeier, Kung-Chi Chang, Ashleigh Frewen, Padraig Taaffe, Joke C. Grans, Haoyu Xiong, Brian A. Benz, Suzanne Russo, Dawid Nowak, Stephen Hearn, Allen Yu, John E. Wilkinson, Frank Rigo, David L. Spector

**Author notes:** Correspondence: Sarah D. Diermeier or David L. Spector.

## Abstract

Long non-coding RNAs (lncRNAs) are an emerging class of regulatory molecules that have been shown to play important roles in tumorigenesis and cancer progression. Here, we studied the recently identified lncRNA *Mammary Tumor Associated RNA 20* (*MaTAR20*) in mammary cancer progression. A CRISPR/Cas9 knockout of *MaTAR20* in the metastatic 4T1 cell line led to reduced cancer cell proliferation and increased cell surface adhesion compared to control cells. Consistent with these knockout results antisense oligonucleotide (ASO) mediated knockdown of *MaTAR20* resulted in reduced growth and invasion in 4T1 cells, and in primary mammary tumor organoids derived from the MMTV-PyMT mouse model of breast cancer. Injection of *MaTAR20*-specific ASOs subcutaneously into tumor bearing MMTV-PyMT mice resulted in smaller and highly necrotic tumors in comparison to mice injected with a scrambled control ASO. To investigate the molecular mechanism by which *MaTAR20* acts to advance mammary tumor progression, we applied a combination of RNA-sequencing and RNA-pulldown coupled to DNA-sequencing. These analyses demonstrated that the nuclear retained lncRNA is associated with several essential cancer signaling pathways such as VEGF signaling. In particular, *MaTAR20* directly binds to and regulates the expression of *Tnfsf15*. Our results indicate that *MaTAR20* is an important driver of mammary tumor progression and represents a promising new therapeutic target.

## Introduction

Breast cancer (BC) is estimated to account for 30% of new cancer diagnoses and 15% of cancer-related deaths in women in 2021 (1). While currently available chemo- and targeted therapies have led to improved overall survival rates, declines of mortality have slowed over the past decade compared to other types of cancer (1). Metastatic disease in particular is the main cause for BC related mortality (1), indicating the need for innovative approaches to target the metastatic cascade.

Recent studies highlight the potential of long non-coding RNAs (lncRNAs) as new therapeutic targets in cancer (2–13). Many lncRNAs are expressed in a tissue- and cancer-specific manner (14, 15) and several previous studies have supported a role for lncRNAs as drivers of tumorigenesis, tumor growth and invasion (reviewed in (16) and (17)). A promising new approach of targeting lncRNAs to reduce mammary tumor growth and metastasis *in vivo* are nucleic acid based therapies (reviewed in (17)). Initial success in targeting oncogenic lncRNAs was shown using locked nucleic acids (LNAs) to target *BCAR4* (5) or antisense oligonucleotides (ASOs) for *Malat1* (6, 18).

We previously identified 30 *<u>Ma</u>mmary <u>T</u>umor <u>A</u>ssociated <u>R</u>NAs* (*MaTARs*) (7). These lncRNAs are over-expressed in mouse models of BC and in human breast tumor tissue compared to normal mammary epithelial cells (7). ASO-mediated knockdown of various *MaTARs* in primary mammary tumor cells resulted in a tumor cell - specific reduction of cell growth (7). Recently, we demonstrated that *MaTAR25* expression contributes to BC progression via regulation of the *Tensin1* gene (13). Here, we investigated the potential of *MaTAR20* (also known as *Gm13387*, *RP23-132N23.1*, *ENSMUSG00000087028*) as a new therapeutic target in BC, as well as the molecular mechanism by which *MaTAR20* acts. ASO-mediated knockdown or CRISPR/Cas9 knockout of *MaTAR20* results in reduced cell proliferation and migration. Further, ASO knockdown of *MaTAR20 in vivo* leads to delayed tumor onset, decreased tumor size, increased tumor necrosis and reduced metastatic burden. To further study the potential of this lncRNAs as a therapeutic target, we generated two loss-of-function and one gain-of-function models of the human orthologue, *hMaTAR20*. Our *in vitro* and *in vivo* characterization of these models suggest that the lncRNA is indeed a new driver of breast cancer cell proliferation and migration. Based on our investigation of the molecular mechanism of *MaTAR20* function, the observed phenotype is likely the result of reduced tumor vascularization upon reduction in the level of *MaTAR20*. *MaTAR20* regulates genes involved in tumor angiogenesis, such as increasing the expression of the *Vascular Endothelial Growth Factor A and B* (*Vegf-A*, *Vegf-B*) genes and it directly binds to the *Tnfsf15* locus to repress its expression during tumor progression.

## Materials and Methods

### Murine cell culture

Murine 4T1, Py2T, G0771 and D2A cells were cultured in DMEM supplemented with 10% FBS and 1% penicillin/streptomycin. All cells were grown in a cell culture incubator at 37 °C and 5% CO2. ASOs were delivered to the cells via free uptake immediately after seeding the cells. For proliferation assays, cells were seeded in 24-well plates and trypsinized at individual time points. Cell counts were determined manually using a hemocytometer. Cell cycle analysis was performed using a BrdU FITC kit (BD) as described (13).

### Organoid culture

Murine mammary tumor organoids were generated and cultured as previously described (7, 19). Briefly, organoids were generated from nulliparous MMTV-PyMT tumors, mixed with growth factor-reduced Matrigel and grown in DMEM/F12 medium supplemented with 1x ITS (insulin, transferrin, and sodium selenite), 1% penicillin/streptomycin and murine FGF2 (2.5 nM). For ASO-mediated knockdown experiments, organoids were seeded at a density of 5 organoids/µl and plated as 80 µl domes in 24-well dishes. ASOs were added directly (free uptake) to the culture medium 15-20 min after the organoids were plated and in fresh medium at day 3. Organoids were incubated for a total of 6 days. ASO sequences are provided in Supplementary Table S1. For visualization purposes and quantification of organoid branching, images were acquired using an Axio-Observer Live Cell inverted microscope (Zeiss).

### Scanning electron microscopy (SEM)

SEM on 2D cells was performed as described previously (20). Briefly, cells cultured on glass coverslips were fixed in 1.6% glutaraldehyde in PBS for 30 minutes and then dehydrated in a graded ethanol series. Coverslips were critical point dried (CPD) using a Samidri 795 CPD device (Tousimis), mounted on aluminum stubs with carbon tabs (Electron Microscopy Sciences). Samples were coated with gold using an Emitech K550X Sputter Coater. Samples were imaged with a Hitachi S3500 SEM operated at 5 kV.

For SEM of 3D organoids, Matrigel containing organoid samples were removed from multi-well tissue culture dishes (using a 3.5 mL disposable transfer pipette and placed in 45 mL of cold PBS (4°C) in 50 mL capacity polypropylene centrifuge tubes (VWR Scientific). Matrigel was washed away by 3-4 rounds of suspension in cold PBS and centrifugation using a refrigerated centrifuge set to 1,000 pm, 20 min, 4°C (Beckman Coulter Allegra X15r). After the final wash to remove Matrigel, organoids were transferred to 1.6 mL microcentrifuge tubes and fixed in 1% glutaraldehyde in PBS at 4°C, for 30 minutes, rinsed in distilled water by centrifugation and resuspension and then post-fixed in 1% aqueous osmium tetroxide (Electron Microscopy Sciences) for 30 minutes at RT. After osmium fixation, organoids became dense relative to solutions and a centrifuge was not needed for further handling. Organoids were dehydrated in a graded ethanol series and then immersed in 50% hexamethyldisilazane (HMDS, Electron Microscopy Sciences) in 100% ethanol for 5 min and 100% HMDS for 10 min. Organoids in 0.3 mL of HMDS were aspirated using a transfer pipette and the solution and suspended organoids were deposited to 12 mm diameter SEM stubs coated with carbon adhesive tabs (Ted Pella Inc) and allowed to air dry. Samples were coated with gold using an Emitech K550X Sputter Coater and imaged with a Hitachi S3500 SEM operated at 5 kV.

### RNA extraction and qRT-PCR

Total RNA was isolated from cells or organoids using TRIzol. DNase I treatment was performed for 15 min at RT to remove contaminating DNA. cDNA synthesis was carried out using the TaqMan Reverse Transcription kit (Life Technologies) and random hexamers according to the manufacturer’s instructions. Quantitative Real time PCR (qRT-PCR) was performed using the Power SYBR Green Master Mix (Life Technologies). Cycling conditions were as follows: 15 min at 95 °C followed by 40 cycles of 15 sec at 94 °C, 30 sec at 60°C. *Peptidylprolyl isomerase B* (*cyclophilin B*) was used as an endogenous control in mouse and *RPL13* in human to normalize each sample and relative expression results were calculated using the 2^-ΔΔCt^ method. A list of primers used is provided in Supplementary Table S1.

### Subcellular localization

For single-molecule FISH, cells were seeded onto acid-washed glass coverslips and fixed in 4% paraformaldehyde when reaching 50% confluency. RNA-FISH was carried out using the Affymetrix View ISH Cell Assay Kit and custom probes (Thermo Fisher) according to the manufacturer’s instructions. For human tissue, Pantomics BRC481 tissue microarrays were used. The slides were imaged on a LSM 710/780 confocal microscope (Zeiss). Cell fractionation assays were performed as described in (13).

### CRISPR/Cas promoter deletion

To generate a genetic knockout, two sgRNAs targeting the promoter region were combined, creating a deletion. A sgRNA targeting the *Renilla* luciferase gene was used as non-targeting control. All sgRNAs were cloned into a lentiCRISPR_V2 plasmid (Addgene #52961) also encoding Cas9 and delivered to the cells using lentiviral transduction as described (21). Stable integrants were selected using puromycin selection and single cell sorted using a FACS Aria (SORP) Cell Sorter. Each single cell clone was propagated and analyzed by genomic PCR and qRT-PCR to select for homozygous knockout clones. Sequences for sgRNAs are provided in Supplementary Table S1.

### Invasion assay

Invasion assays were performed as described previously in (7). Briefly, a Cultrex® 96 well BME Cell Invasion Assay (Trevigen) was used. Cells were starved in FBS-free culture medium, then harvested and seeded at a density of 5 x 10^4^ cells/well into the invasion chamber. ASOs were added to growth medium containing 10% FBS. The plate was incubated at 37 °C for 24 h and the assay was performed according to the manufacturer’s instructions. The fluorescence was measured with a SpectraMax i3 Multi- Mode Detection Platform (Molecular Devices) using the 480/520 nm filter set. Each sample was measured in triplicate.

### Rescue Assay

*MaTAR20* isoforms were amplified by PCR using Phusion High-Fidelity DNA Polymerase (NEB) following the manufacturer’s instructions. *MaTAR20* isoform 1 was cloned using Xhol and EcoRI-HF overhangs. The digested PCR product was ligated into a pCMV6-Entry plasmid (Origene) with T4 DNA ligase. Isoforms 2 and 3 were amplified by PCR, gel extracted and treated with T4 PNK. The pCMV6-Entry plasmid was digested using EcoRV-HF and Eco53kl followed by dephosphorylation using Quick CIP. Isoforms 2 and 3 were ligated into the pCMV6-Entry backbone as described above. Ligations were transformed into competent DH5α cells using heat shock. Transformants were selected for using Kanamycin at 25 µg/mL and validated using Sanger sequencing.

For ectopic expression of *MaTAR20*, knockout cells were seeded into 96-well plates at a density of 5,000 cells/well. The plates were incubated overnight (16-18 h) at 37 °C with 5% CO2. DNA was transfected using Lipofectamine 2000 (Thermo Fisher) in serum-free DMEM medium at a ratio of 100 ng DNA : 0.2 µL Lipofectamine. After 6 h, the medium was replaced with 100 µL/well of fresh complete medium. Cells were fixed for each time point using 100 µL/well of staining solution (150 µg/mL saponin, 4 µg/mL DAPI, 0.5% PFA in PBS) for 30 minutes with gentle shaking at RT, then stored at 4 °C .The cells were counted using a Cytation5 automated cell counting system.

RNA was isolated 24 h after transfection. The medium was removed and 50 µL of TRIzol was added to each well of a 96-well plate. RNA was extracted according to the manufacturer’s instructions. For each sample, 1 µg of RNA was treated with ezDNase and reverse transcribed to cDNA using SuperScript IV VILO following the manufacturer’s instructions (Thermo Fisher). For RT-qPCR testing of the *MaTAR20* rescue, PowerUp SYBR Green Master Mix (Thermo Fisher) was used. All RT-qPCRs were run on a QuantStudioTM 3 Real-Time PCR Machine (Applied Biosystems).

### *hMaTAR20* CRISPR/Cas9 knockout cell lines

T47D cells were cultured in RPMI-1640 medium supplemented with 10% FBS, 0.2 Unit/mL human insulin and 1% penicillin/streptomycin. A sgRNA targeting the mouse *MaTAR20* gene was used as non-targeting control (M888). All sgRNAs were cloned into a lentiCRISPR_V2 plasmid (Addgene #52961) also encoding Cas9 and delivered to the cells using lentiviral transduction as described (21). Stable integrants were selected using puromycin selection and single cell sorted using a FACS Aria (SORP) Cell Sorter. Each single cell clone was propagated and analyzed by genomic PCR and qRT-PCR to select for homozygous knockout clones (19bp deletion; hg38 chr9:137295224-137295246).

### CRISPRi

MDA-MB231-LM2 (LM2) cells (22) were cultured in DMEM supplemented with 10% FBS and 1% penicillin/streptomycin. We used lentiviral transduction to integrate dCas9 (pHAGE EF1α dCas9-KRAB, Addgene # 50919) as described (21). A sgRNA targeting the *Renilla* luciferase gene was used as non-targeting control. sgRNAs targeting the promoter region of *hMaTAR20* (hg38 chr9:137295979-137295998 for -277; chr9: 137296113-137296128 for -390) were cloned into pLenti SpBsmBI Hygro plasmid (Addgene #62205) and delivered to the MDA-MB231-LM2-dCas9 cells using lentiviral transduction. Stable integrants were selected using hygromycin selection. For *in vivo* experiments 400,000 cells were xenografted into the L4 mammary fat pad of Nu/J mice (n=10 each for *Ren* control and *hMaTAR20* KD cells).

### CRISPRa

MCF10A cells were cultured in a media consisting of DMEM/F12 + GlutaMAX + 5% Horse Serum + 20 ng/ml EGF + 0.5 mg/ml Hydrocortisone + 100 ng/ml Cholera Toxin + 0.28 Units/ml Insulin. For CRISPR activation (CRISPRa) two plasmids carrying the components of the tagging system SUperNova (SunTag) were integrated into MCF10A cells by lentiviral transduction (23). The utilized plasmids were obtained from Addgene: pHRdSV40-scFv-GCN4-sfGFP-VP64-GB1-NLS (#60904) and pLV-sgCDKN1B#2 BFP (#60903). All sgRNAs were cloned into the plasmid backbone pLenti SpBsmBI Hygro (Addgene #62205), and stable integrants were selected using hygromycin selection.

### Proliferation assay

The proliferation assay was performed by DAPI staining of the cell nuclei and automatically counted by a CytationTM 5 Cell Imaging Reader (BioTek). Briefly, cells were trypsinized and counted manually with a hemocytometer. One thousand cells per well were seeded into 10 wells of four 96-well plates. Cells were fixed and stained at 24 h, 48 h, 72 h, and 96 h post seeding with 100 μl staining solution (PBS, 150 ng/ml Saponin, 8 ng/ml DAPI, 0,5% PFA) per well for 30 minutes.

### Scratch assay

Scratch assays were performed following the protocol according to (24). Cells were trypsinized, seeded into a 24-well plate (or 6-well plates for MCF10A) and grown to full confluence. The cell monolayer was scratched using a 200 µL pipette tip to create a straight wound, followed by washing off floating cells with medium. Images were taken directly after scratching (T0) and after 24 h culturing (T24), and the rate of wound closure was calculated using Image J.

### Animal experiments

Animal experiments were carried out in the CSHL Animal Shared Resource, in accordance with IACUC approved procedures. MMTV-PyMT mice (25) were obtained from Dr. Mikala Egeblad (CSHL). Tumors and normal mammary glands were extracted immediately after euthanizing the animal and processed to generate primary cells, organoids or tissue sections. For *in vivo* ASO injections, female MMTV-PyMT mice were divided into two cohorts, being treated either with a *MaTAR20*-specific ASO or a scrambled control ASO at ∼8-10 weeks of age (after formation of palpable tumors). MOE ASOs were injected subcutaneously three times per week (n=4 for scASO, n=5 for *MaTAR20* ASO). cET ASOs were injected twice per week, both at 50 mg/kg per dose (n=7 for scASO, n=7 for *MaTAR20* ASO). Tumors were measured twice per week throughout the treatment course. At the end of the experiment, mice were euthanized, and primary tumors and lungs were fixed in 4% PFA and embedded in paraffin for histo- pathological analysis, or snap-frozen for RNA extractions. FFPE blocks were sectioned and stained with hematoxylin and eosin (H&E). Slides were scanned and analyzed using the Aperio ImageScope software. Immunohistochemistry was performed using the following antibodies: Tnfsf15 (Bioss, BS-5092R), Anti-Ki67 (AbCam, ab15580), Cleaved Caspase-3 (Cell Signaling, 9661), VegfB (Invitrogen, PA5-80220), VegfA (AbCam, ab1316). The resulting images were analyzed using QuPath software v0.2.3(26).

### RNA-seq

RNA quality was assessed on an Agilent 2100 Bioanalyzer. Libraries were prepared on samples with RIN ≥9 using the Illumina TruSeq sample prep kit v2 and sequenced on an Illumina NextSeq instrument. Data was analyzed as previously described (7). Briefly, the quality of FASTQ files was assessed using FastQC (http://www.bioinformatics.babraham.ac.uk/projects/fastqc/). Reads were mapped to GRCm38 using STAR (27), and the reads per gene record were counted using HTSeq-count (28). Differential gene expression was performed with DESeq2 (29), and an adjusted p-value of < 0.05 was set as threshold for statistical significance. KEGG pathway and GO term enrichment and was carried out using the R/Bioconductor packages GAGE (30) and Pathview (31).

### ChIRP-seq

ChIRP-seq was carried out as previously described (13). Briefly, 20 million cells were harvested and fixed in 1% glutaraldehyde solution. ChIRP was performed using two individual tiling pools of biotinylated oligonucleotide probes. A probe pool targeting mouse *Ppib* was used as negative control. A list of ChIRP-seq probes is provided in Supplementary Table S1. ChIRP-Seq libraries were generated using the Illumina TruSeq ChIP Library Preparation Kit and sequenced on an Illumina NextSeq instrument. ChIRP-seq data quality was assessed using FastQC (http://www.bioinformatics.babraham.ac.uk/projects/fastqc/) and paired-end reads were mapped to GRCm38 using Bowtie2 (32). ChIRP-seq analysis was performed using HOMER (33). For RT-qPCRs testing the ChIRP cDNA, 5 ng cDNA and 5 µM of primers in 10 µL was used (reagents and cycling conditions as described under “Rescue assay”).

### Data access

RNA-seq and ChIRP-seq data have been deposited at Gene Expression Omnibus (GEO) under accession number GSE171085.

## Results

### *MaTAR20* is a tumor-specific, nuclear-retained lncRNA

*MaTAR20* is an intergenic lncRNA gene located on mouse chromosome 2. According to GENCODE vM18, three isoforms of the *MaTAR20* transcript exist, comprising 2,646, 659 and 638 nt (Figure 1A). Our computational analysis of ENCODE expression data revealed that the lncRNA is absent or expressed at low levels in normal mammary glands and other normal tissues, but highly expressed (FPKM >200) in mammary tumors derived from the transgenic MMTV-PyMT mouse model of luminal BC (Figure 1B). Furthermore, *MaTAR20* levels correlate with tumor size in the MMTV-PyMT model (Supplementary Figure S1A).

**Figure 1:**
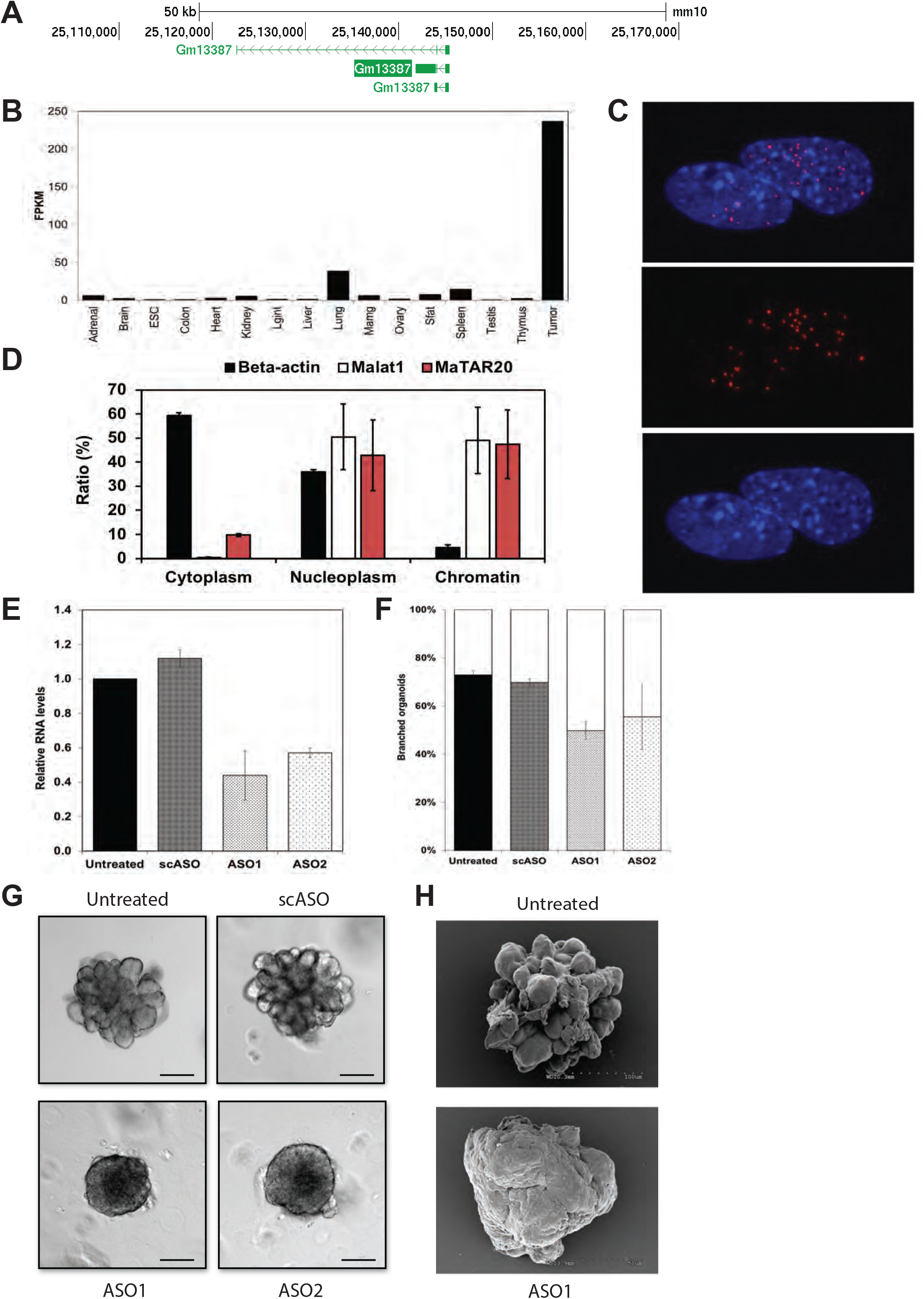
*MaTAR20* is a tumor-specific, nuclear-retained lncRNA. **A:** Schematic overview of the *MaTAR20*/*Gm13387* locus (UCSC Browser, mm10) GENCODE vM18 annotation in green. **B:** Abundance (FPKM) of *MaTAR20* in ENCODE mouse tissue and HER2+ mammary tumors. **C:** Single molecule RNA FISH of MaTAR20 in primary MMTV-PyMT cells. Top panel: merge, middle panel: MaTAR20 (red), bottom panel: DAPI (blue). **D:** Cellular fractionation assay followed by qRT-PCR. **E:** ASO-mediated knockdown of MaTAR20 in primary MMTV-PyMT organoids. Two specific ASOs were tested for MaTAR20, and a scASO was used as negative control. Bars denote the mean of at least two biological replicates +/- SD. **F:** Quantification of organoid branching upon MaTAR20 knockdown. Bars denote the mean of at least three biological replicates +/- SD. **G:** Exemplarily images of organoids quantified in F. **H:** Exemplarily SEM micrograph of organoids quantified in F.

To examine the localization of *MaTAR20* we performed single molecule RNA-FISH in primary MMTV-PyMT cells (Figure 1C). Our results indicate that *MaTAR20* is indeed an abundant lncRNA in primary mammary tumor cells, and predominantly localizes to the nucleus, averaging 10-40 foci per nucleus. We further confirmed that *MaTAR20* is a nuclear-retained transcript by subcellular fractionation assays, which revealed an enrichment in both the nucleoplasm and the chromatin fractions (Figure 1D), a localization pattern comparable to the lncRNA *Malat1*. Our initial characterization indicates that *MaTAR20* is a tumor-specific, nuclear-enriched lncRNA that may play a role in tumor progression based on its expression profile.

### Knockdown of *MaTAR20* leads to reduced organoid branching

To determine if *MaTAR20* is a driver of mammary tumor cell growth, we performed knockdown experiments *ex vivo*, in MMTV-PyMT derived tumor organoids (6,7,34) (Figure 1E-H). Organoids are an excellent model system to study cancer biology and to test new therapeutic treatments (for review, see (35)). Wildtype/untreated organoids exhibit a branched morphology consistent with collective cell migration (Fig 1G-H) and/or proliferation (for review, see (36). We achieved *MaTAR20* knockdown efficiencies of 50- 60% using 2’-O-methoxyethyl (MOE) ASOs in 3D mammary tumor organoids by free uptake (Figure 1E). Treatment of MMTV-PyMT tumor organoids with two independent MOE ASOs targeting *MaTAR20* lead to an up to 40% reduction of organoid branching compared to untreated organoids, or organoids treated with a scrambled control MOE ASO (scASO) (Figure 1F-H). Based on these and our previous results in primary mammary epithelial cells (7), we conclude that *MaTAR20* enhances both the proliferative and migratory potential of mammary tumor cells.

### Promoter deletion of *MaTAR20* results in a less aggressive phenotype

To independently validate our ASO-mediated knockdown results, and to develop a model system that enables us to study the molecular mechanism of *MaTAR20* in detail, we generated *MaTAR20* loss-of-function cells using the CRISPR/Cas9 system. First, we tested a panel of mouse mammary tumor cell lines to identify a suitable model system. Of the tested cell lines, the metastatic 4T1 line (37) showed the highest *MaTAR20* expression level (Supplementary Figure S2A). Next, we established promoter deletion of *MaTAR20* by stably integrating Cas9 and two guide RNAs (gRNAs): the first gRNA was designed to bind upstream of the *MaTAR20* transcription start site (at -736) and the second gRNA binding within the first exon of *MaTAR20* (+12 bp) (Figure 2A), resulting in promoter deletion of 748 bp. A gRNA targeting the *Renilla* luciferase gene was used to generate negative control cells. All edited cell populations were single cell sorted to generate monoclonal cell lines. Two *MaTAR20* knockout clones (KO1, KO2) and two negative control clones (Ren1, Ren2) were analyzed further. The KO cell lines showed a ∼80-85% reduction of *MaTAR20* expression compared to control clones (Figure 2B). We used four different primer pairs to measure *MaTAR20* levels, one amplifying all three isoforms of *MaTAR20* (PP1, Figure 2B), and three isoform-specific primer pairs (PP2, PP3, PP4; Supplementary Figure S2B). Our results indicate that all three isoforms of *MaTAR20* were reduced to comparable levels in the KO cells.

**Figure 2:**
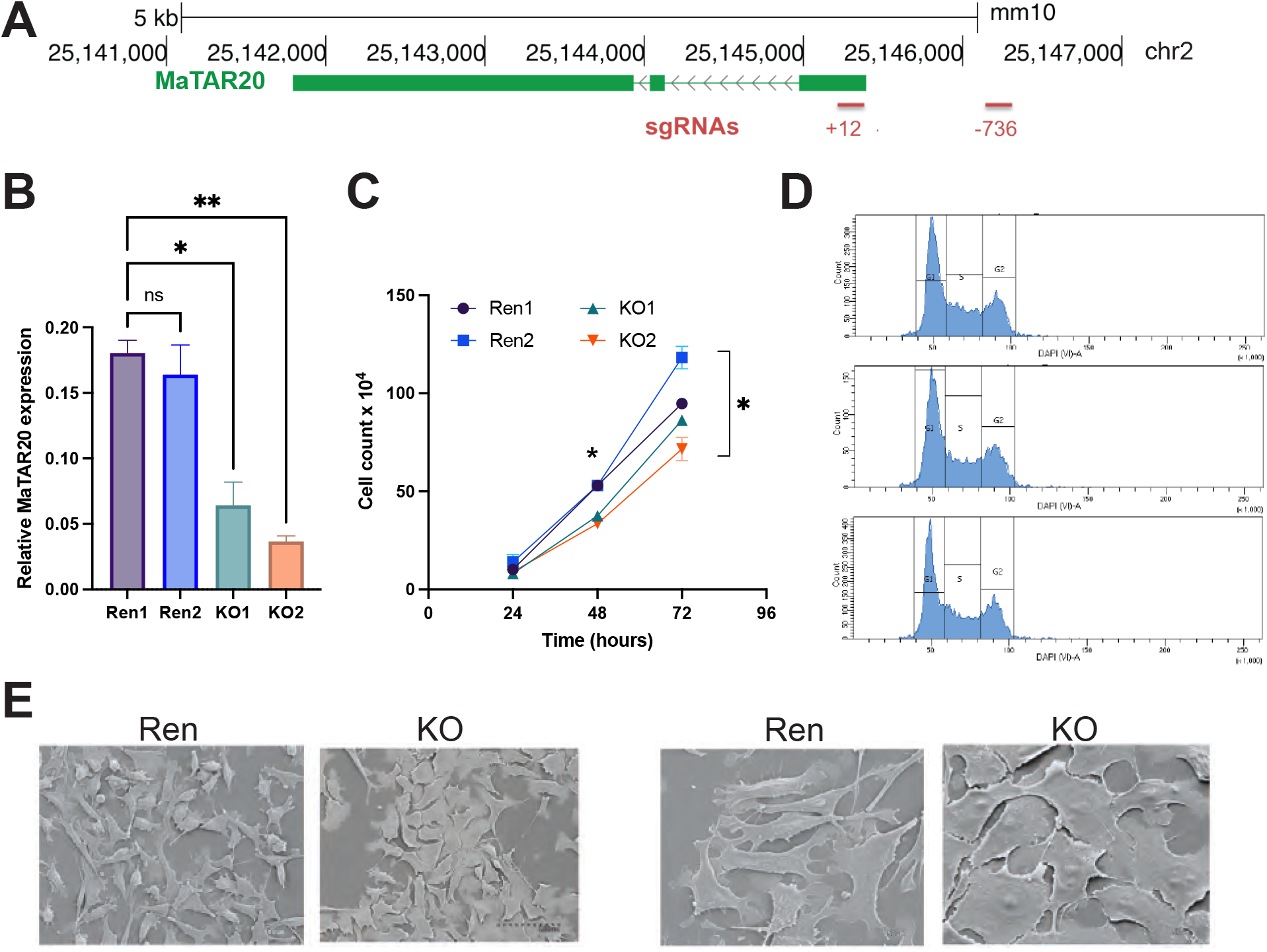
*MaTAR20* KO cells show reduced proliferation and cell adhesion. Statistical significance was determined with a two-way ANOVA; * p < 0.05, ** p < 0.01. **A:** Schematic overview of the knockout strategy. Guide RNAs (gRNAs) are indicated in red, +/- indicate genomic position relative to the *MaTAR20* transcription start site. **B**: qRT-PCR to determine relative *MaTAR20* expression in *MaTAR20* promoter deletion cell lines. KO = promoter deletion of 748 bp (combination of gRNAs “+12” and “-736”), Ren = negative control, integration of Cas9 and a gRNA targeting *Renilla* luciferase. Bars denote the mean of at least six biological replicates +/- SEM. **C**: Cell proliferation assay. Number of cells determined after 24, 48 and 72h. Data points denote the mean of at least two biological replicates +/- SEM. **D:** Cell cycle profiles of NST-DAPI stained cells, comparing Ren control cells to KO1 and KO2. **E:** Scanning electron micrograph of control (Ren) or CRISPR/Cas9-modified cells (KO). Scale as indicated.

Similar to our ASO-mediated knockdown experiments in primary mammary tumor cells (7), we observed a 30-40% reduction of cell proliferation in the KO cell lines (Figure 2C). This phenotype was not caused by differences in cell cycle profiles (Figure 2D) and did not correlate with Cas9 expression (Supplementary Figure S2C). Ectopic expression of *MaTAR20* in KO cells was able to rescue the cell proliferation phenotype, restoring growth rates to levels comparable to Ren control cells (Supplementary Figure S3A-B). This finding also indicates that the *MaTAR20* transcript is responsible for the observed phenotype, rather than the genomic locus itself playing a regulatory role and that *MaTAR20* plays a role in *trans*.

While culturing the *MaTAR20* KO cells, we noticed a difference in cell adhesion between the KO clones and the control cell lines. The KO cells were particularly difficult to dislodge from cell culture flasks, leading to increased (2-4 fold) trypsinization times required to detach the KO cells. Further examination of the KO and Ren control cells using scanning electron microscopy revealed that reduced *MaTAR20* levels may lead to stronger surface attachment of the cells, accompanied by more organized, sheet-like structures (Figure 2E). In summary, promoter deletion of *MaTAR20* leads to reduced tumor cell growth and migration compared to control cells, similar to primary cells treated with *MaTAR20* ASOs (Figure 1).

### The human orthologue, *hMaTAR20,* drives breast cancer cell growth

We previously identified the human orthologue of *MaTAR20*, *hMaTAR20*, by synteny as *RP13-122B23.8 (ENSG00000260996*), located on chromosome 9 (7). To investigate whether *hMaTAR20* drives breast cancer cell growth and therefore may hold clinical significance, we first performed computational analyses of *hMaTAR20* expression of TCGA data. The lncRNA is overexpressed 2-fold across all subtypes of breast cancer compared to adjacent normal breast tissue, with highest levels in estrogen receptor positive (ER+) breast cancer (Figure 3A). RNA-FISH on a tissue microarray further confirmed a 2-fold up-regulation of *hMaTAR20* in breast cancer compared to corresponding normal tissue (Figure 3B). Further stratifying the breast cancer tissue revealed highest levels of *hMaTAR20* in grade II-III tumors (Supplementary Figure S4A), following a similar trend of increased expression in later stage tumors as observed in mouse (Supplementary Figure S1).

**Figure 3:**
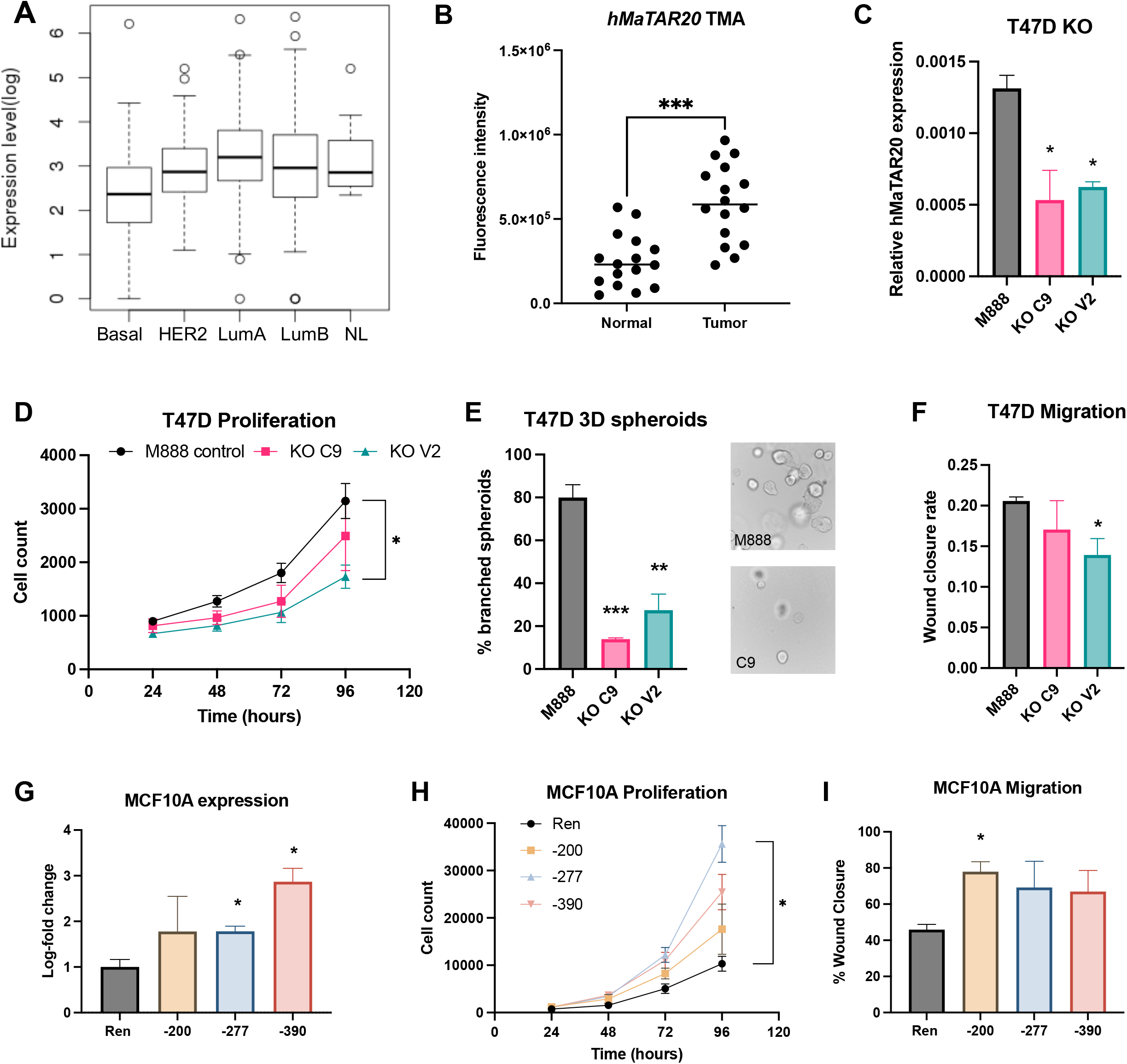
Human *MaTAR20* (*hMaTAR20*) drives cell proliferation and migration. Bars denote the mean of three biological replicates +/- SEM. * p < 0.05, ** p < 0.01, *** p < 0.001. **A:** Analysis of *hMaTAR20* expression in TCGA data, by subtype. LumA = luminal A, LumB = luminal B, NL = normal-like breast cancer. **B:** RNA-FISH of *hMaTAR20* on a tissue microarray (TMA), comparing expression in breast cancer to corresponding normal breast tissue. **C:** qRT-PCR to determine relative *hMaTAR20* expression in CRISPR/Cas9 edited T47D cells. M888: control cell line with non-targeting sgRNA. KO C9 and KO V2: individual single cell sorted colonies of T47D cells with a deletion in exon 1. **D:** Cell proliferation assay comparing growth curves of the M888 negative control to the two *hMaTAR20* KO clones. **E:** Left: quantification of T47D spheroid branching (left). Right: exemplary images of the negative control M888 (top) and the KO C2 clone (bottom). For each biological replicate, n = 100 spheroids were analyzed (n = 300, total). **F:** Quantification of T47D scratch assay. **G:** qRT-PCR to determine relative *hMaTAR20* expression in MCF10A cells. The endogenous *hMaTAR20* gene was up-regulated using CRISPRa using three specific sgRNAs: -200, -270 and -390 (numbers denote the distance from the TSS). A non- targeting sgRNA for *Renilla* luciferase served as a negative control (Ren). **H:** Cell proliferation assay comparing growth curves of the Ren negative control to the three *hMaTAR20* CRISPRa cell lines. **I:** Quantification of MCF10A scratch assay.

We performed an expression screen in several breast cancer cell lines, which further confirmed that ER+ cell lines show increased levels of *hMaTAR20* (Supplementary Figure S4B), with T47D showing the highest levels. Therefore, we generated a CRISPR/Cas9 driven deletion in exon 1 in T47D cells, which resulted in stable reduction of *hMaTAR20* levels (Figure 3C) in two single cell clone derived cell lines (KO C9 and KO V2). Lower *hMaTAR20* levels lead to reduced cell proliferation in T47D KO cells compared to control cells grown in 2D culture and reduced branching in spheroid culture, with more pronounced effects in the 3D assay (Figure 3D-E). We also observed reduced 2D migration in *hMaTAR20* KO cells (Figure 3F). Like *MaTAR20* in mouse, *hMaTAR20* showed predominantly nuclear localization (Supplementary Figure S4C), suggesting the lncRNA functions similarly in both organisms.

While *hMaTAR20* expression is highest in ER+ breast cancer, the lncRNA is also overexpressed in all other types of breast cancer compared to adjacent normal tissue. To study the impact of *hMaTAR20* in an ER- model, we generated a CRISPR interference (CRISPRi) driven transcriptional repression model of *hMaTAR20* in the aggressive basal cell line MDA-MB-231-LM2 (22)(Supplementary Figure S4D). Similar to the T47D KO cells, we observed reduced cell proliferation, spheroid branching and cell migration in the MDA-MB-231-LM2 CRISPRi model (Supplementary Figure S4E-G).

Next, we tested whether increasing *hMaTAR20* levels in the pre-malignant cell line MCF10A may lead to an increase in proliferative and migratory potential. MCF10A cells express *hMaTAR20* at very low levels compared to T47D or MDA-MB-231-LM2 cells (Figure 3B). Using CRISPR activation (CRISPRa), we up-regulated endogenous *hMaTAR20* expression using three different sgRNAs (Figure 3G). We observed increased cell proliferation and viability as well as increased migration compared to control cells (Figure 3H-I). In summary, our results show that *hMaTAR20* is a driver of cell proliferation and migration in human breast cancer cells, recapitulating the phenotype we observed with *MaTAR20* in the mouse system.

### cET ASO shows improved knockdown efficiency *in vitro*

To determine the effect of *MaTAR20* reduction on tumor progression, and to investigate whether ASOs are a viable strategy to systemically reduce *MaTAR20*, we performed *in vivo* experiments in transgenic mouse models of breast cancer. First, we designed a constrained ethyl (cET) ASO targeting *MaTAR20*. In comparison to MOE ASOs (Figure 1), cET ASOs show increased potency *in vivo* (38) and are the chemistry of choice for tissues less sensitive to ASO modulation with several clinical trials currently ongoing (39).

Before introducing the cET ASO into animals, we tested its knockdown (KD) efficiency *in vitro*. Concentrations as low as 250 nM were sufficient to achieve >60% knockdown of *MaTAR20* within 24 h in 4T1 cells (Figure 4A), and we observed KD efficiencies of >95% using 500 nM after 72 h (Figure 4B). These results compare favorably to concentrations of 4-5 µM of MOE ASOs achieving ∼50% KD in organoids (Figure 1) and ∼70% KD in primary cells (7). Knockdown of *MaTAR20* using the cET ASO resulted in significantly reduced cell proliferation and invasion (Figure 4C-D), however, it is possible that the difference in proliferation impacts on the readout of the invasion assay as well. These results agree with our observations using MOE ASOs (Figure 1) and *MaTAR20* promoter deletions in mouse (Figure 2) as well as our findings for *hMaTAR20* (Figure 3).

**Figure 4:**
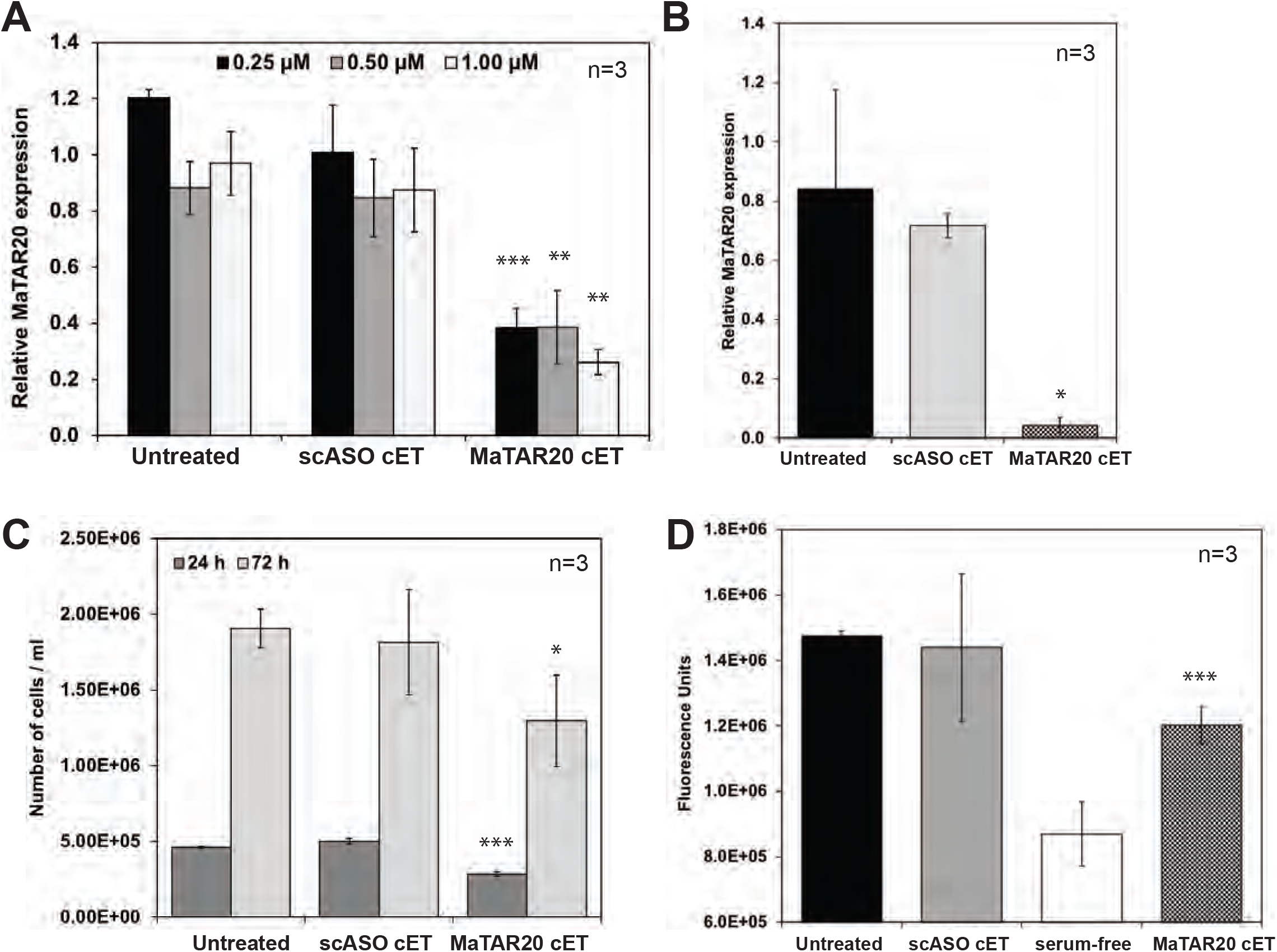
cET ASO-mediated knockdown of *MaTAR20* in 4T1 cells lead to reduced cell proliferation and invasion. Bars denote the mean of three biological replicates +/- SEM. Statistical significance was determined with a two-tailed Student’s t-test; * p < 0.05, ** p < 0.01, *** p < 0.001. **A:** qRT-PCR to determine relative *MaTAR20* expression. 24 h post treatments with cET ASOs (concentrations as indicated). **B:** qRT-PCR to determine relative *MaTAR20* expression. 72 h post treatments with 0.5 µM cET ASOs. **C:** Proliferation assay. Cells were counted 24 h and 72 h post treatment with 0.5 µM cET ASOs. **D:** Invasion assay. Fluorescence units 24 h post treatment with 0.5 µM cET ASOs.

### Knockdown of *MaTAR20 in vivo* leads to a reduction in tumor growth and induces tumor necrosis

To further investigate the potential of cET ASOs targeting *MaTAR20* in mammary tumors, we injected 100 mg/kg/week of either *MaTAR20* cET or a negative control ASO (scrambled = scASO cET) into MMTV-PyMT (C57BL/6) mice after palpable tumor formation (n=7 for scASO, n=8 for *MaTAR20* ASO; Figure 5A). We achieved an 80% reduction of *MaTAR20* in the tumors on average, further highlighting the potency of the cET chemistry (Figure 5B). We initiated ASO injections on animals with comparable average tumor burden in both cohorts and observed a 30% overall reduced tumor burden in *MaTAR20* cET treated mice over the course of eight weeks (Figure 5C). While mice in the *MaTAR20* cET group eventually went on to develop tumors, tumor onset was delayed in comparison to the scASO cET group. Furthermore, when comparing the growth curves of similarly sized tumors that developed at the same time in both cohorts, tumors in the *MaTAR20* cET group grew slower compared to scASO cET treated mice, resulting in a difference of 75% after 8 weeks (Figure 5D). After all mice reached the study endpoint, we carried out hematoxylin and eosin (H&E) staining of the fixed tumors to investigate potential histo-pathological differences. Notably, tumors in the *MaTAR20* cET treated group showed severe necrosis compared to the control group (Figure 5E).

**Figure 5:**
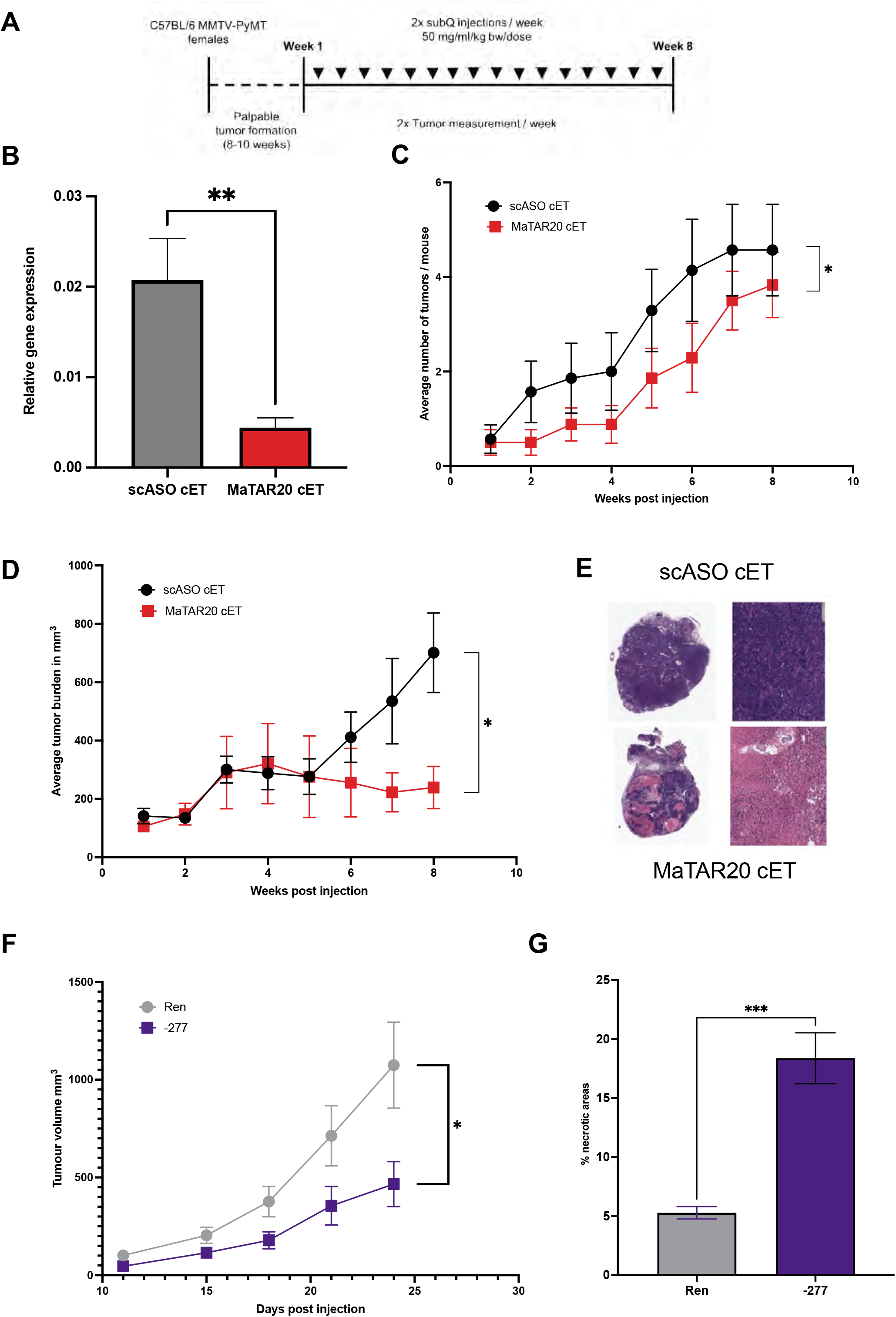
cET ASO-mediated knockdown of *MaTAR20 in vivo* lead to reduced tumor growth and metastatic burden. Statistical significance was determined with a two-tailed Student’s t-test; * p < 0.05, ** p < 0.01, *** p < 0.001. **A:** MMTV-PyMT (C57/Bl6) mice were treated for 8 weeks with cET ASOs (100 mg/kg/week), either scASO (cET) or *MaTAR20* cET1. Bars / lines denote the mean of biological replicates +/- SEM. N= 33 tumors from 7 mice for scASO (cET); n = 26 tumors from 8 mice for *MaTAR20* cET1. **B:** qRT-PCR to determine relative *MaTAR20* expression in tumors. **C:** Average number of tumors per mouse in each treatment group. **D:** Average tumor burden in mm^3^ per week in each treatment group. **E:** Hematoxylin and Eosin staining of tumor sections. Top left: scASO (cET) treated tumor (whole tumor). Top right: scASO (cET) treated tumor, 10x. Bottom left: *MaTAR20* cET1 treated tumor (whole tumor). Bottom right: *MaTAR20* cET1 treated tumor, 10x. **F:** Tumor growth curves of MDA-MB-231-LM2 cells were xenografted into nude mice. Ren: a non-targeting negative control (n=10); -277: *hMaTAR20* CRISPRi cell line (n=9). Data points denote the mean of biological replicates +/- SEM. **G:** Quantification of tumor necrosis, based on H&E staining. Bars denote the mean of three biological replicates +/- SEM.

To confirm that the human orthologue of *MaTAR20* acts similarly to the mouse lncRNA in an *in vivo* situation, we xenografted one of our loss-of-function models, the modified MDA-MB-231-LM2 cells into the mammary fat pad of nude mice. We observed reduced tumor growth and increased tumor necrosis (Figure 5F-G), further confirming the functional role of the *hMaTAR20* in tumor progression.

### Knockdown of *MaTAR20 in vivo* leads to reduced metastatic burden

The MMTV-PyMT model is highly aggressive, with a metastasis incidence rate of >80% (25). We examined the lungs in our *in vivo* study post-mortem for macro-metastatic nodules. While 7/7 mice in the scASO group developed at least one macro-metastatic nodule, only 4/7 mice in the *MaTAR20* ASO cohort presented macro-metastatic lesions (Supplementary Figure S5A). We went on to examine the histo-pathology of the four lungs that developed metastasis by H&E staining and discovered that metastases in the *MaTAR20* ASO treated group showed cystic, differentiated nodules compared to the solid masses detected in the scASO cohort (Supplementary Figure S5B-C). We suggest that metastatic nodules in animals that received the *MaTAR20* cET may have developed prior to treatment start, and that the cystic phenotype is a consequence of systemic *MaTAR20* knockdown, impacting already existing metastases. Alternatively, the observed differences could be due to reduced overall tumor burden.

To independently validate our animal studies, we performed a second *in vivo* experiment using *MaTAR20* MOE ASO1 and injected it into MMTV-PyMT (FVB) mice (Supplementary Figure S5D). Similar to the cET experiments in C57/Bl6, we observed reduced tumor growth for the *MaTAR20* ASO group compared to the scASO group from around week 4 of treatment (Supplementary Figure S5E). H&E staining of tumor sections revealed that animals in the *MaTAR20* MOE group also developed more necrotic tumors compared to control animals (Supplementary Figure S5G). In addition, we observed a reduction of macro-metastatic nodules in the lungs of 3/4 mice receiving *MaTAR20* MOE ASO1 (Supplementary Figure S5F), confirming the trend of reduced metastasis upon *MaTAR20* knockdown.

Immunohistochemistry staining of the tumor sections for Ki67 revealed slightly reduced levels in both MOE and cET cohorts. We did not observe a consistent trend when staining for the apoptosis marker cleaved caspase 3, indicating that the tumors likely undergo necrosis rather than programmed cell death (Supplementary Figure S6A-B). In summary, our *in vivo* experiments show that systemic reduction of *MaTAR20* using ASOs leads to delayed mammary tumor onset, reduced tumor growth, increased tumor necrosis and reduced metastasis.

### *MaTAR20* expression level correlates with VEGF expression

We set out to identify the molecular mechanism by which *MaTAR20* impacts tumor cell growth and invasion. As our RNA-FISH and subcellular fractionation assays indicated that *MaTAR20* is a nuclear-retained lncRNA (Figure 1), we hypothesized that it may act by impacting gene expression. First, we performed a computational co-expression analysis using lncRNA2function (40). Similar approaches have been previously applied successfully to identify the molecular role of other lncRNAs (41). Notably, we observed a strong enrichment for pathways involved in angiogenesis, with the top 7 most enriched pathways being related to vascular endothelial growth factor (VEGF) signaling (Figure 6A), including “Signaling by VEGF”, “VEGF hypoxia and angiogenesis” and “Sorafenib Pharmacodynamics”, a multi-kinase inhibitor that has been shown to act on VEGF receptor (VEGFR) (42). Due to this striking ontology enrichment, we set out to experimentally assess if *MaTAR20* and VEGF expression correlate. We determined by qRT-PCR that Vegf-A and Vegf-B are reduced in tumors of MMTV-PyMT mice that received the *MaTAR20* cET ASO compared to the scASO control group (Figure 6B). Immunohistochemistry staining of tumor sections confirmed this trend on the protein level (Supplementary Figure S6C-D). We also assayed CD31 staining in the tumors to probe for differences in angiogenesis and observed reduced CD31+ cells in *MaTAR20* ASO treated animals (Supplementary Figure S6E), which may point to a reduced number of blood vessels being formed upon *MaTAR20* loss.

**Figure 6:**
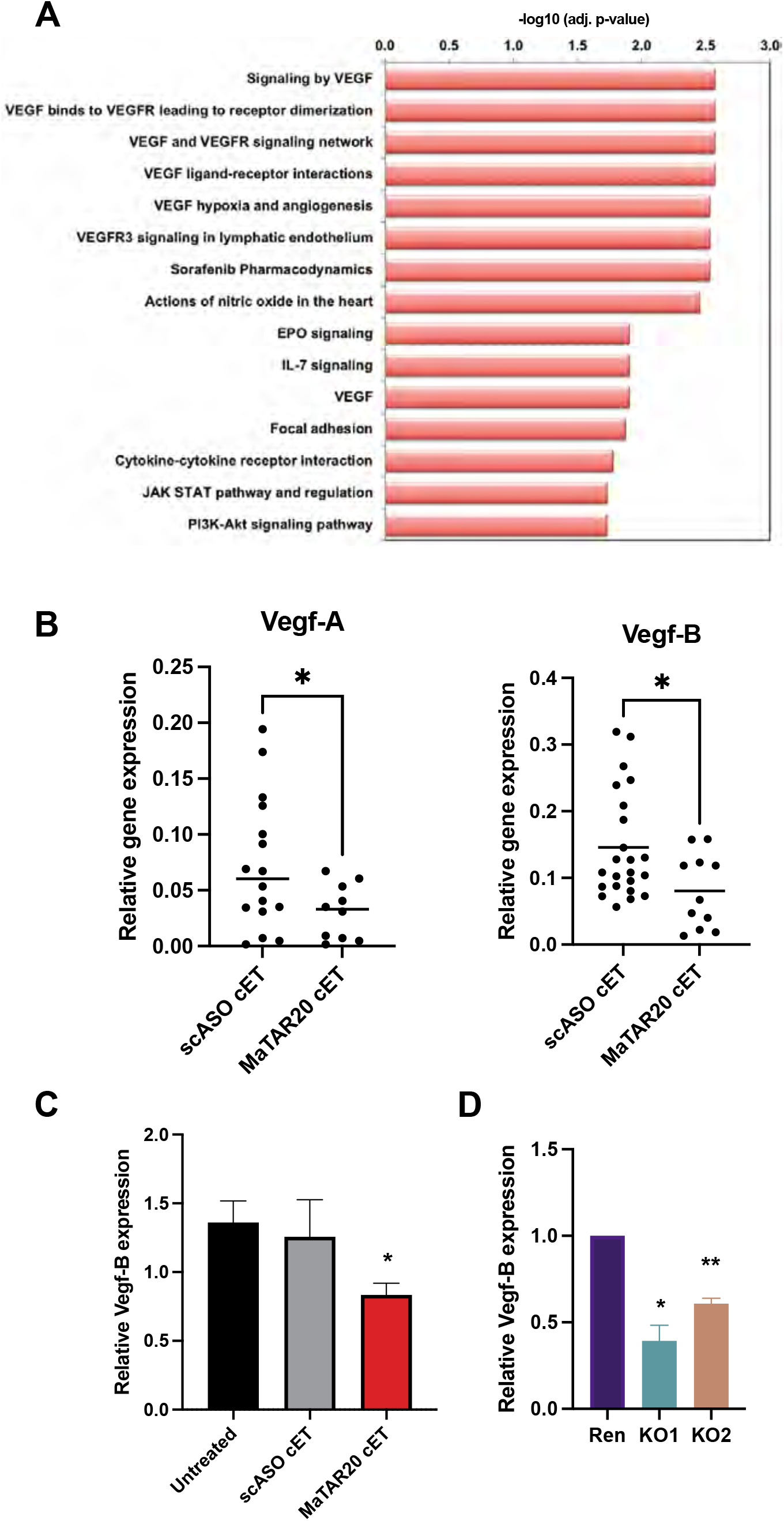
*MaTAR20* impacts VEGF expression. Statistical significance was determined with a two-tailed Student’s t-test; * p < 0.05, ** p < 0.01. **A:** Co-expression analysis to identify *MaTAR20* associated pathways. The 15 most significant pathways are shown. **B:** qRT-PCR to determine the relative Vegf-A and Vegf-B expression in tumors. MMTV- PyMT mice were treated for 8 weeks with cET ASOs (100 mg/kg/week), either scASO (cET) or *MaTAR20* cET1. **C:** qRT-PCR to determine the relative Vegf-B expression in 4T1 cells. 72 h post treatments with 0.5 µM cET ASOs. Bars denote the mean of three biological replicates +/- SEM. **D:** qRT-PCR to determine the relative Vegf-B expression in *MaTAR20* KO cells. Bars denote the mean of two biological replicates +/- SEM.

We used RNA extracted from whole tumors for the qRT-PCR validations described above. While MMTV-PyMT tumors generally contain a high percentage of cancer cells (Figure 5E, Supplementary Figure S5E), qRT-PCR from whole tumors cannot distinguish between expression changes in tumor or stromal cells. To investigate whether VEGF expression was reduced in the same cell type as *MaTAR20*, we measured VEGF expression in 4T1 cells with successful *MaTAR20* KD (Figure 4A/B). Indeed, we observed a 35% reduction of Vegf-B (Figure 6C), while Vegf-A was not expressed at levels sufficient for reliable qPCR detection in these cells (Ct > 30). We also tested Vegf-B expression in our three knockout cell lines (Figure 2), and observed a statistically significant, 40-60% down-regulation of Vegf-B compared to control cells (Figure 6D). Upon ectopic expression of *MaTAR20* in the KO cells, Vegf-B levels were up-regulated (Supplementary Figure S7A). Based on convergent results obtained from four independent experiments (ASO KD *in vitro*, *in vivo*, CRISPR/Cas9 promoter deletions and rescue assays), we conclude that down-regulation of *MaTAR20* correlates with reduced VEGF expression levels and may be associated with reduced tumor angiogenesis.

### *MaTAR20* loss leads to alterations in cancer signaling and adhesion pathways

To identify how *MaTAR20* impacts Vegf signaling, we further investigated the molecular mechanism by which the lncRNA acts. First, we performed differential RNA-seq, comparing *MaTAR20* KO to control cells. A total of 223 genes were differentially expressed (p < 0.05), with 129 genes down- and 94 genes up-regulated in the KO cells (Figure 7A, Supplementary Figure S7B, Supplementary Table S2). Interestingly, one of the most up-regulated genes was *Vascular endothelial growth inhibitor* (*Vegi*), also known as *Tumor necrosis factor (TNF)-like cytokine 1A (TL1A)/TNF superfamily member 15* (*Tnfsf15*). In agreement with our co-expression analysis (Figure 6A), we observed an overall dysregulation of the VEGF signaling pathway and of hypoxia-inducible factor 1 (HIF-1) signaling, another pathway essential for angiogenesis (43) (Supplementary Table S3). In addition, pathways involved in cancer signaling such as the MAPK, PI3K-Akt and

**Figure 7:**
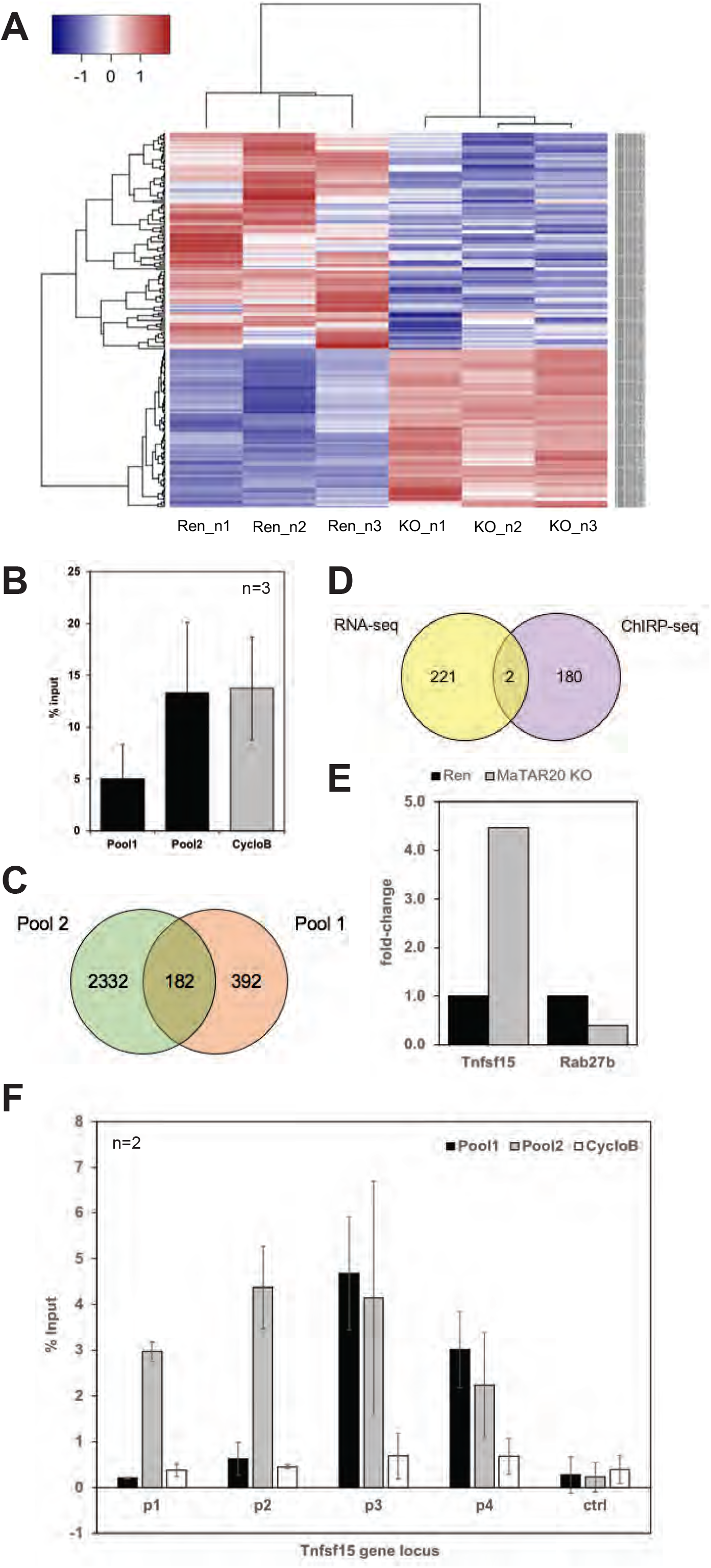
RNA-seq and ChIRP-seq identify *Tnfsf15* as a direct *MaTAR20* target gene. **A:** RNA-seq. Heatmap of significantly differentially expressed genes comparing *MaTAR20* KO and Ren cells. N1-3 indicate three independent biological replicates. **B:** ChIRP-seq. qRT-PCR to determine the efficiency of the ChIRP assay. Both *MaTAR20* probe pools enrich *MaTAR20*, with Pool 2 being more efficient. “Cyclo B” indicates a separate pull-down experiment with oligonucleotides specific for the *cyclophilin B* mRNA, which was used as a specificity control. Percentage of MaTAR20 or CycloB pull-down in relation to input is shown. Bars denote the average of at least 3 replicates +/- SD. **C:** ChIRP-seq. Venn diagram illustrating the overlap of ChIRP-seq performed with Pool1 vs Pool2 probes. Genes also identified in the negative control (Cyclo B) pull-down were removed from the target lists. 183 genes overlap between both ChIRP probe pools, with Pool1 enriching more genes than Pool 2 overall. All ChIRP experiments were performed in three independent replicates. **D:** Venn diagram illustrating the overlap of the 182 target genes identified by ChIRP-seq and the 223 differentially expressed genes according to RNA-seq. 2 genes are both direct genomic binding partners of *MaTAR20* and are differentially expressed upon *MaTAR20* loss. **E:** RNA-seq. Expression of Tnfsf15 and Rab27b, comparing the mean log2 fold-change in the two *MaTAR20* KO cell lines (“MaTAR20 KO”) to the mean log2 fold-change of the Ren negative control cell lines (“Control”). **F**: qRT-PCR to validate direct *MaTAR20* binding to the *Tnfsf15* locus. Four tiling primer pairs were designed to two putative *MaTAR20* binding regions in proximity to the *Tnfsf15* gene. A negative control region, located between p1/p2 and p3/p4, shows no *MaTAR20* enrichment. CycloB probes control for unspecific binding at all tested sites. Bars denote the mean of at least two biological replicates +/- SD.

TNF axes were altered, as were several cell adhesion pathways, including “Focal adhesion” and “Tight junctions” (Supplementary Table S3). Several of these pathways were previously identified in our co-expression analysis as well (Figure 6A). Differential expression of important cell signaling genes likely contributes to the altered phenotypes we observed upon *MaTAR20* loss, such as reduced organoid branching and tumor cell growth. Changes in cell adhesion pathways may explain the enhanced adhesion phenotype observed in *MaTAR20* KO cells (Figure 2E), along with reduced migration and metastatic burden upon KD of the lncRNA.

### *MaTAR20* regulates gene expression by direct binding to *Tnfsf15*

RNA-seq identifies all differentially expressed genes upon *MaTAR20* loss, both effects due to direct binding and regulation of the gene by *MaTAR20*, as well as any secondary or tertiary downstream effects. To hone in on direct targets of *MaTAR20*, we performed Chromatin Isolation by RNA Purification (ChIRP, (44)) using two separate pools of nine tiling oligonucleotides each. Both pools were able to enrich *MaTAR20*, with pool 2 showing overall better pull-down performance (Figure 7B). The difference in enrichment could be due to one or several oligonucleotides in pool 1 being obstructed from binding to *MaTAR20*, caused by secondary structures or competition with putative protein interaction partners. A negative control pull-down reaction for the mRNA *cyclophilin B* (*CycloB*) was performed as well to control for unspecific binding of nucleic acids to biotinylated oligonucleotides and/or streptavidin beads (Figure 7B).

We performed three independent replicates of ChIRP-seq with both probe pools and *CycloB* negative control probes, and compared genes bound to *MaTAR20* in both pools. The highest genomic peak score in both pools was as expected to the *MaTAR20* locus itself, a similar observation was made in regard to *MaTAR25* lncRNA (13). Both pool 1 and pool 2 were enriched for genes involved in cancer signaling and adhesion pathways that were also identified in our RNA-seq analysis, such as PI3K-Akt and MAPK signaling, “Focal adhesion” and “ECM-receptor interactions” (Supplementary Table S4). The more efficient pool 2 pull-down was also enriched for VEGF and HIF-1 signaling pathways (Supplementary Table S4). On an individual gene level, we assigned the identified peaks to the nearest gene and were able to identify 574 genes bound by *MaTAR20* in pool1 and 2,514 genes in pool 2 (excluding *MaTAR20* itself, as well as all genes that were also found to bind to the negative control *CycloB* probe set) (Supplementary Table S5), with an overlap of 182 target genes bound by *MaTAR20* in both probe pools (Figure 7C, Supplementary Table S5).

Comparing RNA-seq and ChIRP-seq results, two genes were identified to be direct binding partners of *MaTAR20* and also differentially expressed in *MaTAR20* KO cells (Figure 7D): *Tnfsf15* and *Rab27b*. While *Tnfsf15* was significantly up-regulated upon *MaTAR20* loss, *Rab27b* was repressed (Figure 7E). Up-regulation of *Tnfsf15* was confirmed at the protein level in immunohistochemistry assays on tumor sections from *MaTAR20* ASO treated animals (Supplementary Figure S6F). To further establish a direct relationship between *MaTAR20* and *Tnfsf15* expression levels, we performed rescue assays over-expressing ectopic *MaTAR20* in KO cells and observed decreased expression of *Tnfsf15* as compared to the *MaTAR20* KO cells treated with just the bavkbone plasmid (Supplementary Figure S7C).

We performed ChIRP-qPCR at the *Tnfsf15* gene locus to validate direct binding of the lncRNA (Figure 6F). Our *MaTAR20* ChIRP-seq data identified two putative lncRNA binding regions, one about 5 kb downstream and the second about 15 kb downstream of the *Tnfsf15* gene (Supplementary Figure S7D). We designed two primer pairs for each region (p1 and p2 for -15 kb, p3 and p4 for -5 kb), and a primer pair for a negative control region in between the two putative binding sites, at about -10 kb. Cyclo B probes served as a negative control for all tested regions, representing background noise. Our results indicate that both *MaTAR20* probe pools were able to specifically detect the lncRNA at the -5 kb site, while only the more efficient pool2 could also enrich for MaTAR20 at the - 15 kb region. This may indicate higher levels of MaTAR20 binding to the *Tnfsf15* -5 kb site, which likely represents a *Tnfsf15* enhancer element. Overall, our results confirm that *MaTAR20* can regulate gene expression by directly binding to target genes such as *Tnfsf15*.

Based on our findings, we conclude that *MaTAR20* reduces or inhibits the expression of *Tnfsf15* by directly binding to its genomic locus. Tnfsf15 is a cytokine that has been described to inhibit VEGF expression (45) and, more generally, VEGF driven angiogenesis (46). Therefore, we hypothesize that loss of *MaTAR20* leads to reduced tumor angiogenesis (Figure 8). The proposed molecular mechanism agrees with the observed phenotype: delayed tumor onset, smaller tumors and increased necrosis, combined with lower metastatic burden in the lungs.

**Figure 8:**
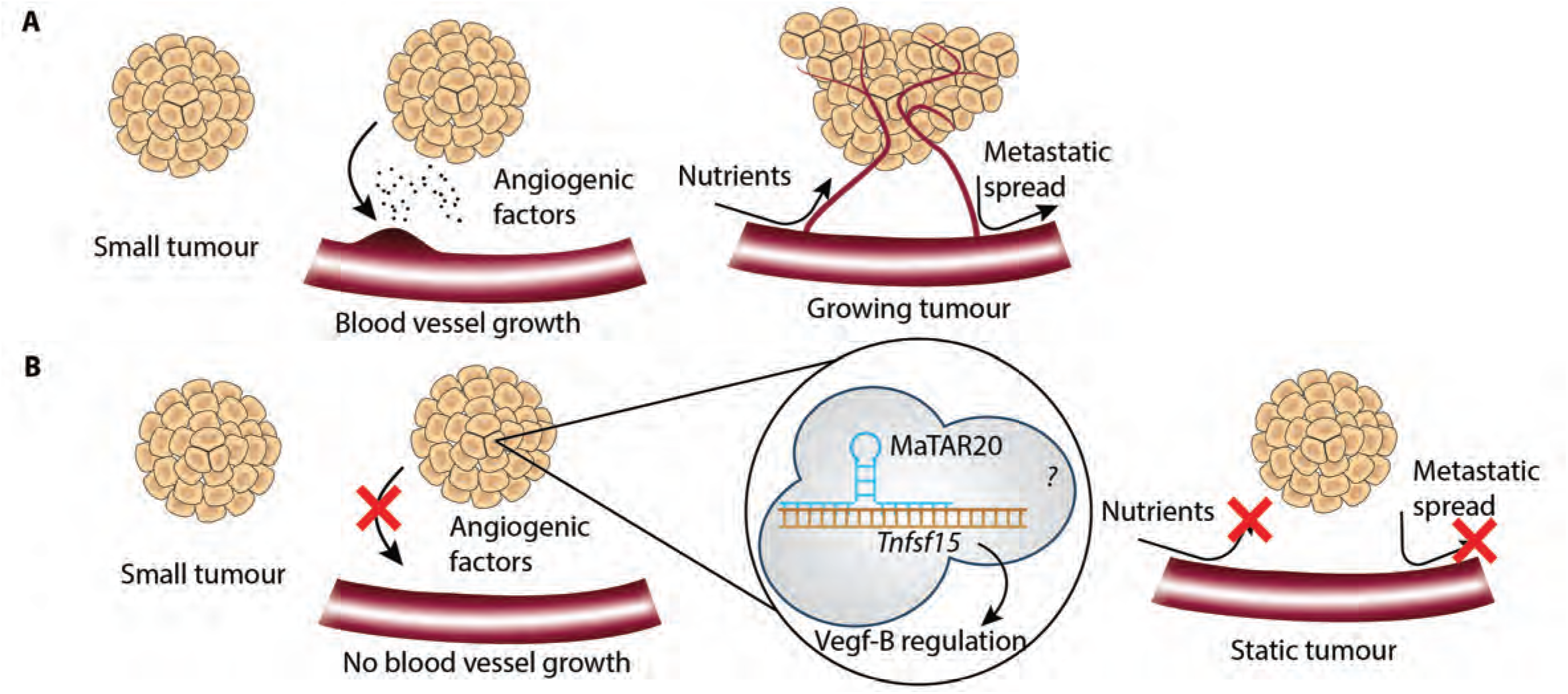
*MaTAR20* knockdown leads to smaller, necrotic tumors and reduced metastasis. **A:** During cancer progression, tumors secret pro-angiogenic factors to recruit blood vessels. Increased vasculature allows for steady tumor growth while also increasing the chances for metastatic spread. **B:** In the absence of *MaTAR20*, tumor growth is slowed down, tumor necrosis increases, and metastasis is reduced. We hypothesize that this could be due to the observed down-regulation of *Vegf-B*, which may in part be caused by direct binding of *MaTAR20* to *Tnfsf15*. Additional players such as protein binding partners may be of importance and will be elucidated further in future studies.

## Discussion

Here, we investigated *MaTAR20*, a lncRNA expressed at high levels in mammary tumors. Loss of *MaTAR20* by either ASO-mediated knockdown or CRISPR/Cas9 genome editing in mammary tumor cells leads to reduced proliferation, invasion and organoid branching *in vitro*. In mouse models of BC, treatment with ASOs targeting *MaTAR20* causes delayed tumor onset, reduced tumor growth and overall reduced metastatic burden to the lungs. We were able to confirm the observed phenotype using different ASO sequences, different ASO chemistries (MOE and cET), and in different mouse background strains (FVB and C57/Bl6). Furthermore, our studies of human *MaTAR20* in two loss-of-function and one gain-of-function models further corroborate that this lncRNA is a driver of BC cell growth and migration and may represent a promising new therapeutic target. We propose that ASOs targeting *MaTAR20* could be a viable new adjunct therapy, as we obtained highly efficient lncRNA knockdown in tumors upon subcutaneous delivery. We did not observe any adverse reactions in other mouse tissues or the animal overall, potentially because *MaTAR20* is expressed at highest levels in the tumor, restricting the effect of *MaTAR20* KD mostly to tumor tissue. Additional detailed studies of the human orthologue *hMaTAR20* (7) will be essential to investigate the potential of the lncRNA as a new therapeutic target in breast cancer. We observed cystic, differentiated metastases in *MaTAR20* ASO-mediated mice, suggesting the treatment may have potential to impact already existing metastases in the lungs. Future investigation of the impact of *MaTAR20* on already existing metastatic nodules will be of particular value, as patients with advanced disease would benefit from a therapeutic approach that is able to attack and/or prevent the formation of metastases.

Our molecular data indicates that the nuclear-retained lncRNA *MaTAR20* drives tumor growth and metastasis by impacting the expression of important cancer signaling pathways and cell adhesion molecules. These findings are reflected in the observed reduction of cell proliferation, migratory potential and formation of cell protrusions in 2D and 3D cell culture systems. One particular pathway that stood out across all our analyses is VEGF signaling and angiogenesis, seemingly centering on Vegf-B in our system. VEGF-B produced by cancer cells has been described to lead to leaky vascular networks, which in turn enables the tumor to invade its surrounding tissue with high efficiency (47). In addition, high VEGF-B levels were associated with poor prognosis (47). Inhibition of tumor-specific VEGF-B has been suggested as an interesting approach to inhibit cancer progression and metastasis, with VEGF-B knockdown leading to reduced pulmonary metastasis in a mouse model of melanoma (47). As *MaTAR20* also leads to reduced lung metastatic nodules, we hypothesize that it may do so by regulating Vegf-B expression. In addition, non-angiogenic functions of VEGF-B have been described previously in the context of invasive BC, which could also play a role here (48).

Our ChIRP assays indicate that *MaTAR20* does not directly bind to the *Vegf-B* gene, but to a number of other angiogenesis-related genes such as *angiopoietin 2*, among many others. Of the 182 genes identified by ChIRP-seq analysis, two also show altered expression in *MaTAR20* KO cells: *Tnfsf15* and *Rab27b*. Tnfsf15 is a cytokine usually expressed in established vasculature but down-regulated in cancer lesions to allow for neovascularization (46,49,50). Vegf and Tnfsf15 serve opposing functions, with carefully balanced expression modulation of both factors playing key roles in vascular and immune homeostasis. We suggest that *MaTAR20* regulates the balance of these proteins by directly binding to the *Tnfsf15* locus and repressing the gene during tumor progression (Figure 8). Reduced Tnfsf15 levels allow the tumor to recruit blood vessels, which is essential for sustained tumor growth and enables intravasation of cancer cells into the circulation, leading to metastasis (Figure 8A). In the absence of *MaTAR20*, we hypothesize that Tnfsf15 expression will be re-activated and neovascularization of the tumor is inhibited, which would agree with our observed reduction in Vegf-A, Vegf-B and CD31+ cells (Figure 8B). In BC, high levels of Tnfsf15 in clinical samples were associated with disease-free survival and overall better prognosis (51). This further solidifies our hypothesis that tumor-specific up-regulation of Tnfsf15 via ASO-mediated *MaTAR20* knockdown could be a promising therapeutic approach. As the lncRNA is expressed in high levels only in cancer lesions, we suggest that consequences of *MaTAR20* KD, including reduced neovascularization, are restricted to tumor tissue as well, representing an attractive new approach of tumor-specific inhibition of angiogenesis.

In addition to regulating angiogenesis, Tnfsf15 has also been described to be involved in dendritic cell maturation and T-cell co-activation (50). Future studies are required to conclude if loss of *MaTAR20* also impacts immune cells in a cancer context, or if the observed phenotype is in fact driven by reduced vascularization. Further investigation is also required into the second identified *MaTAR20* target gene, *Rab27b*. The small secretory GTPase has been described to control vesicle exocytosis and to deliver pro-invasive growth regulators into the tumor microenvironment (52), and has recently been associated with VEGF signaling in cancer as well (53). Rab27b has been reported to promote proliferation and invasiveness of ER+ BC cells *in vitro* and *in vivo*, and has been associated with lymph node metastasis and differentiation grade in ER+ human tumors (54). We observed that loss of *MaTAR20* leads to reduced Rab27b expression, which may further contribute to the decrease in proliferative and migratory potential we observed here. Future studies will elucidate the detailed regulatory mechanism of Rab27b by *MaTAR20*.

## Acknowledgements

We thank members of the Spector and Diermeier labs for critical discussions and advice throughout the course of this study. We thank Dr. Mikala Egeblad (CSHL) for providing MMTV-PyMT mice and Dr. Fred R. Miller (Wayne State University) for providing 4T1 cells. We acknowledge the CSHL Cancer Center Shared Resources (Microscopy, Flow Cytometry, Animal, Histology, and Next-Gen Sequencing) for services and technical expertise (NCI 2P3OCA45508). We thank David Jackson (CSHL) and Zachary Lippman (CSHL) for facilitating access to the SEM instruments. MCF10A and T47D cells used in this study were a generous gift of the Dunbier lab (Department of Biochemistry, University of Otago). We would also like to thank Robert Porteous and the Histology facility at the University of Otago and Bronwyn Carlisle (Department of Biochemistry) for her assistance with the *MaTAR20* model illustration (Figure 8).

---

**Figure S1:**
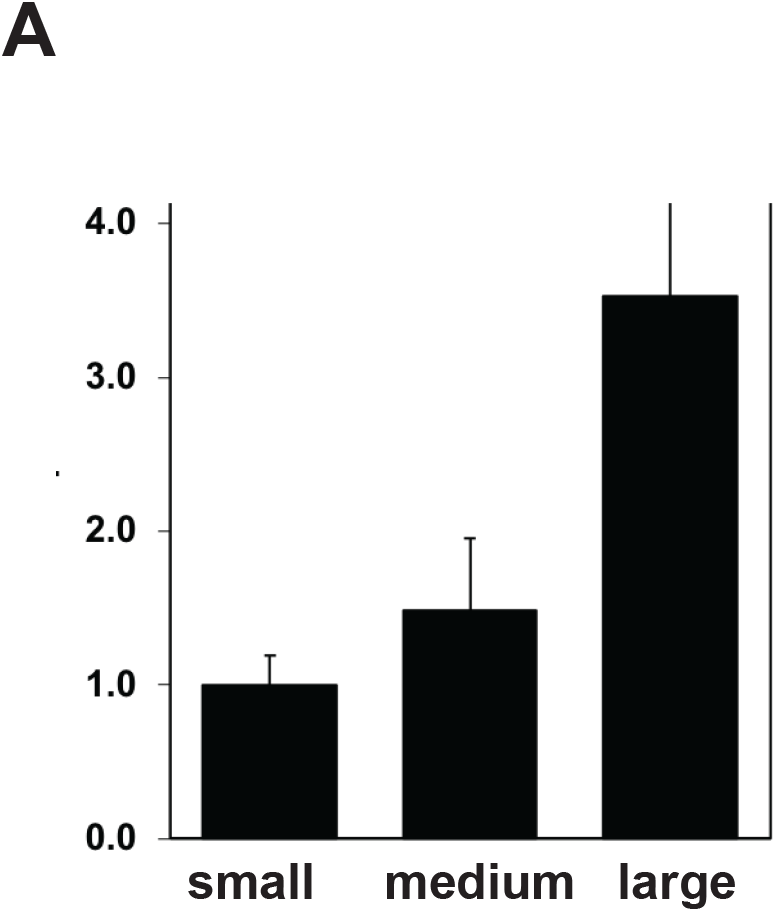
**A:** qRT-PCR to determine relative *MaTAR20* expression in MMTV-PyMT tumors of different size. “Small” tumors: <200 mm^3^, n=7; “medium” tumors: 300 - 1000 mm^3^, n=4; “large” tumors: >1500 mm^3^, n=3.

**Figure S2:**
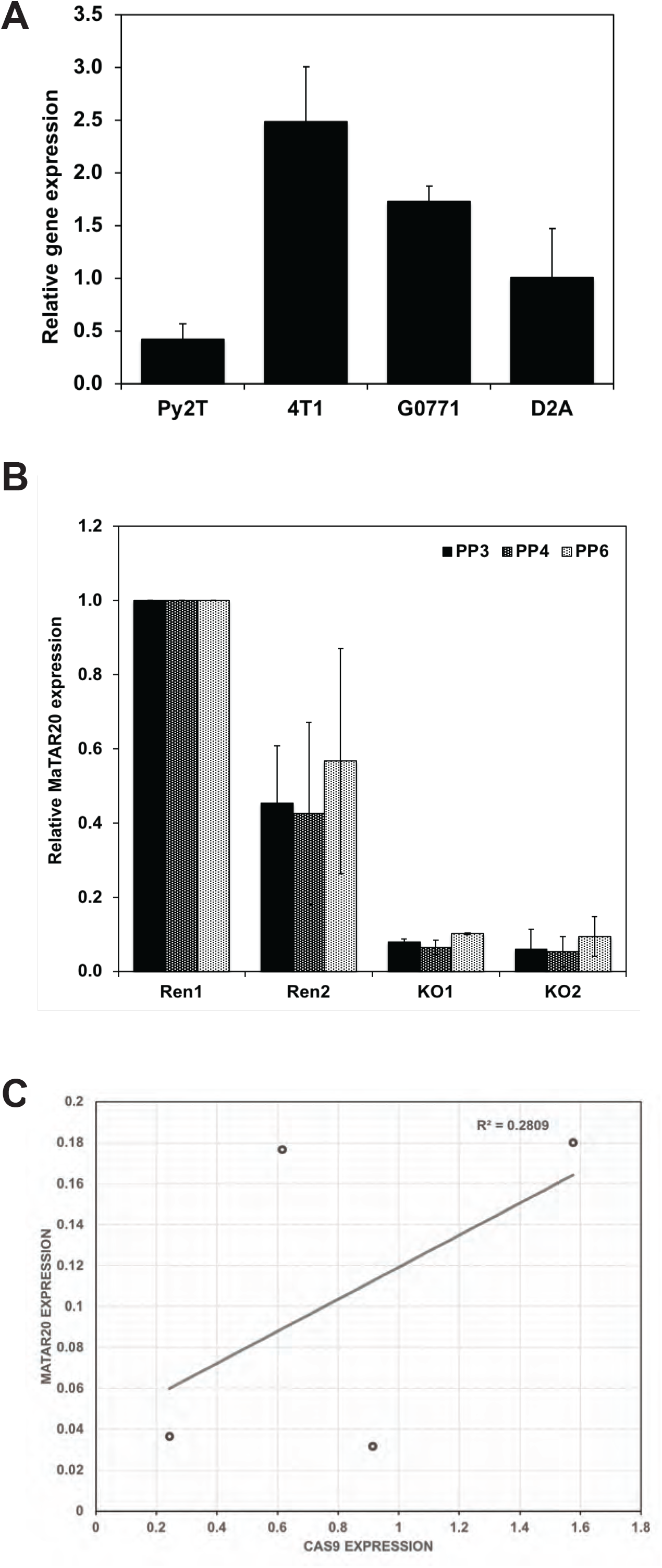
**A:** qRT-PCR to determine the relative *MaTAR20* expression in mouse mammary tumor cell lines. Bars denote the mean of three biological replicates +/- SD. **B:** qRT-PCR to determine the relative expression of individual MaTAR20 isoforms in *MaTAR20* promoter deletion cell lines. KO = promoter deletion of 748 bp (combination of gRNAs “+12” and “-736”), Ren = negative control, integration of Cas9 and a gRNA targeting *Renilla* luciferase. Bars denote the mean of two biological replicates +/- SD. **C:** Correlation of *MaTAR20* expression (all isoforms, based on six biological replicates, Figure 2B) and Cas9 expression, as determined by qRT-PCR in three biological replicates.

**Figure S3:**
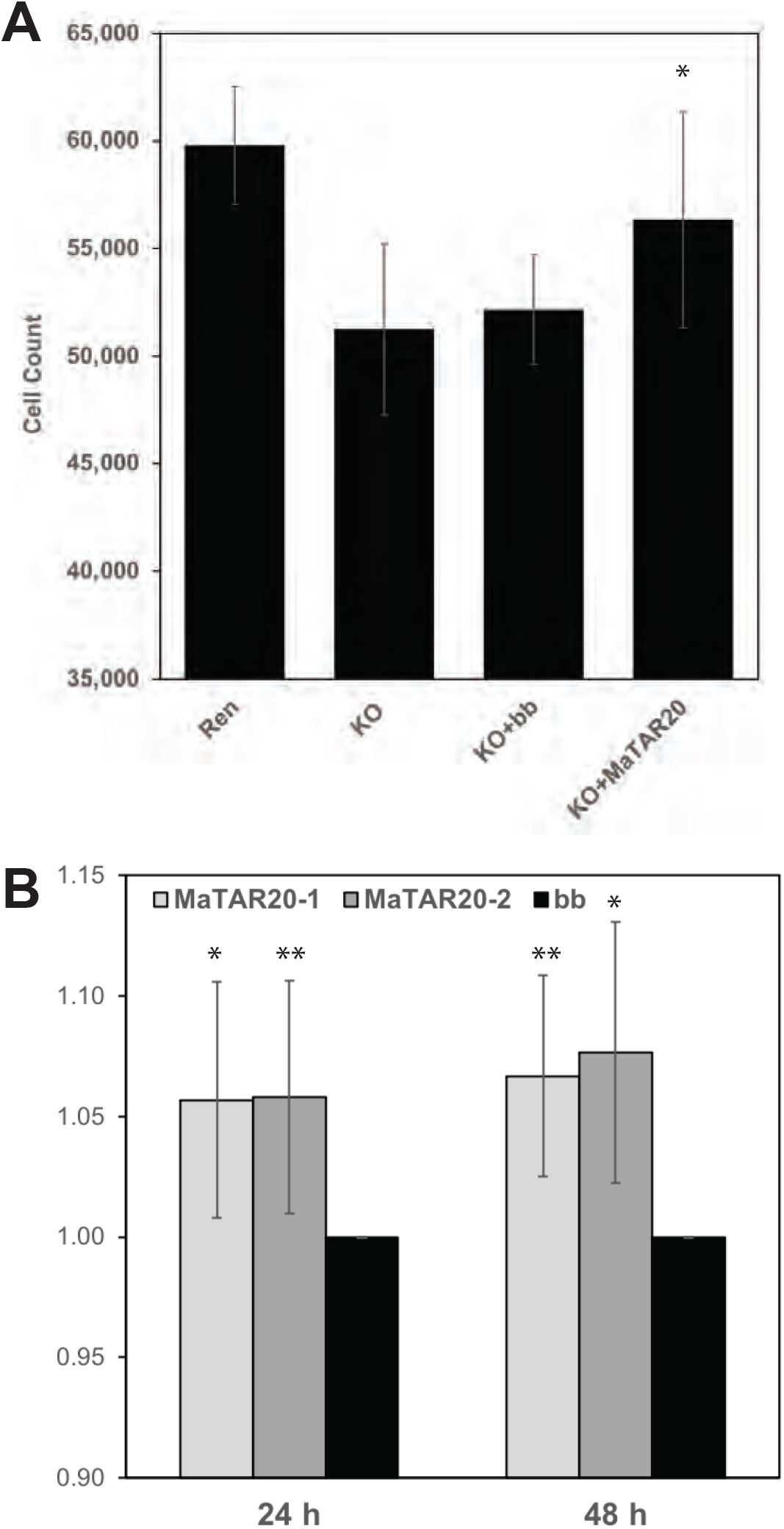
**A:** Cell proliferation assay. Number of cells determined after 48 h. Statistical significance was determined comparing KO cells transfected with a *MaTAR20* Tx1 (KO+MaTAR20) to KO cells transfected with an empty backbone (KO+bb) and untransfected KO cells. Control Ren cells are shown for reference. Error bars denote SD, and a two-tailed Student’s t-test was performed comparing KO+bb to KO+*MaTAR20*; * p < 0.05. **B:** Cell proliferation assay comparing *MaTAR20* isoform 1 and isoform 2 after 24 and 48 h. Cell counts normalized to KO cells transfected with empty vector (bb). Bars denote the mean of at least two biological replicates +/- SD. Statistical significance was determined with a two-tailed Student’s t-test comparing KO+bb to KO+*MaTAR20*; * p < 0.05, ** p < 0.01

**Figure S4:**
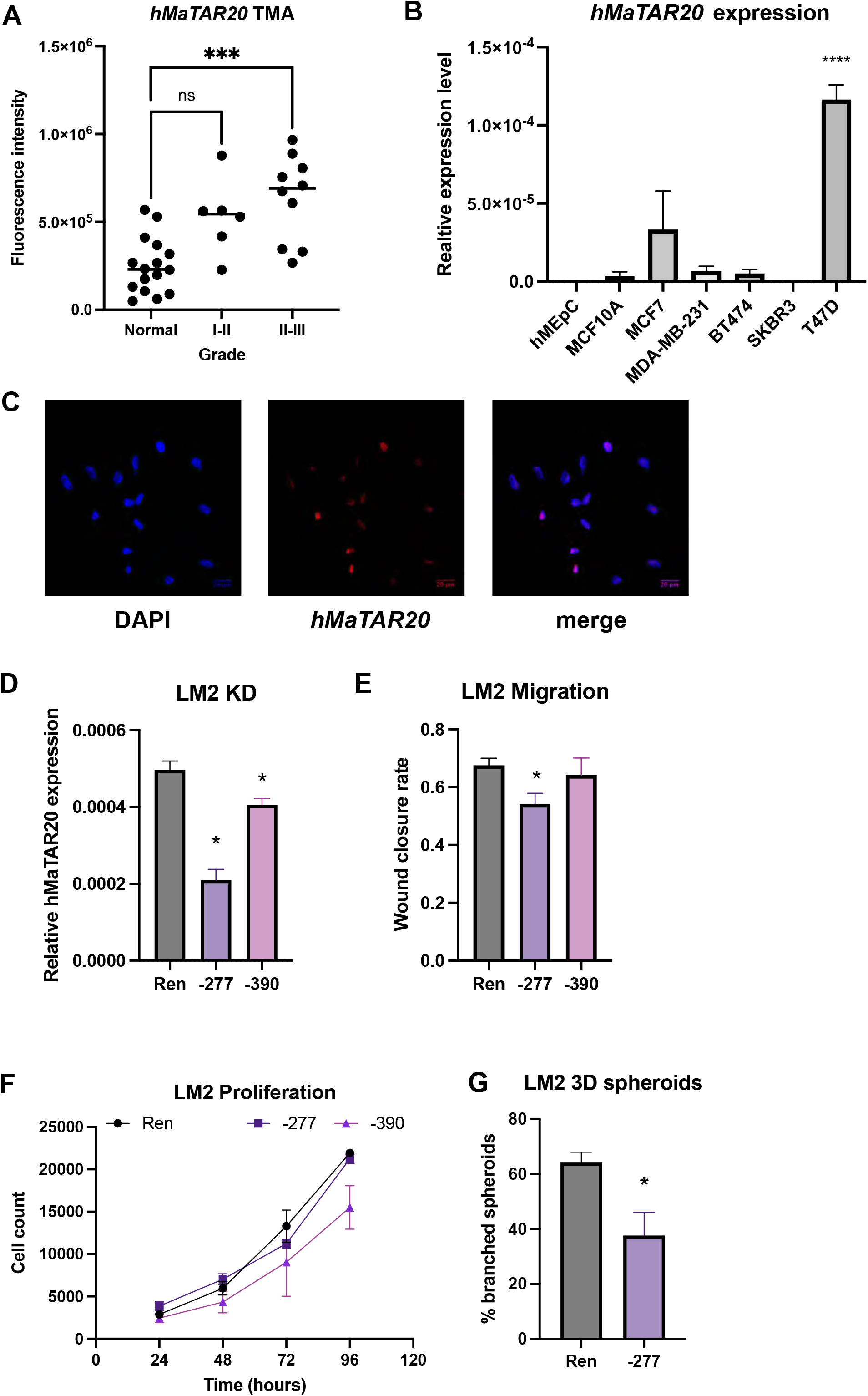
**A:** RNA-FISH of *hMaTAR20* on a tissue microarray (TMA), comparing expression in breast cancer of different grades to corresponding normal breast tissue. Statistical significance was determined with a one-way ANOVA; *** p < 0.001. **B:** qRT-PCR to determine relative *hMaTAR20* expression in a panel of breast cancer cells. Bars denote the mean of three biological replicates +/- SEM. Statistical significance was determined with a one-way ANOVA ;**** p < 0.0001. **C:** RNA-FISH of *hMaTAR20* in MDA-MB-231-LM2 (LM2). **D:** qRT-PCR to determine relative *hMaTAR20* expression in MDA-MB-231-LM2 (LM2) cells. Ren: control cell line with non-targeting sgRNA. -277; -390: CRISPRi *hMaTAR20* knockdown cell lines, numbers denote distance to the TSS. Bars denote the mean of at least three biological replicates +/- SEM. Statistical significance was determined with a two-tailed Student’s t-test; * p < 0.05. **E:** Quantification of LM2 scratch assay. Bars denote the mean of at least three biological replicates +/- SEM. Statistical significance was determined with a two-tailed Student’s t- test; * p < 0.05. **F:** Cell proliferation assay comparing growth curves of the Ren negative control to the two *hMaTAR20* KD cell lines. Data points denote the mean of at least three biological replicates +/- SEM. **G:** Quantification of LM2 spheroid branching. For each biological replicate, n = 100 spheroids were analyzed (n = 300, total). Bars denote the mean of at least three biological replicates +/- SEM. Statistical significance was determined with a two-tailed Student’s t- test; * p < 0.05.

**Figure S5:**
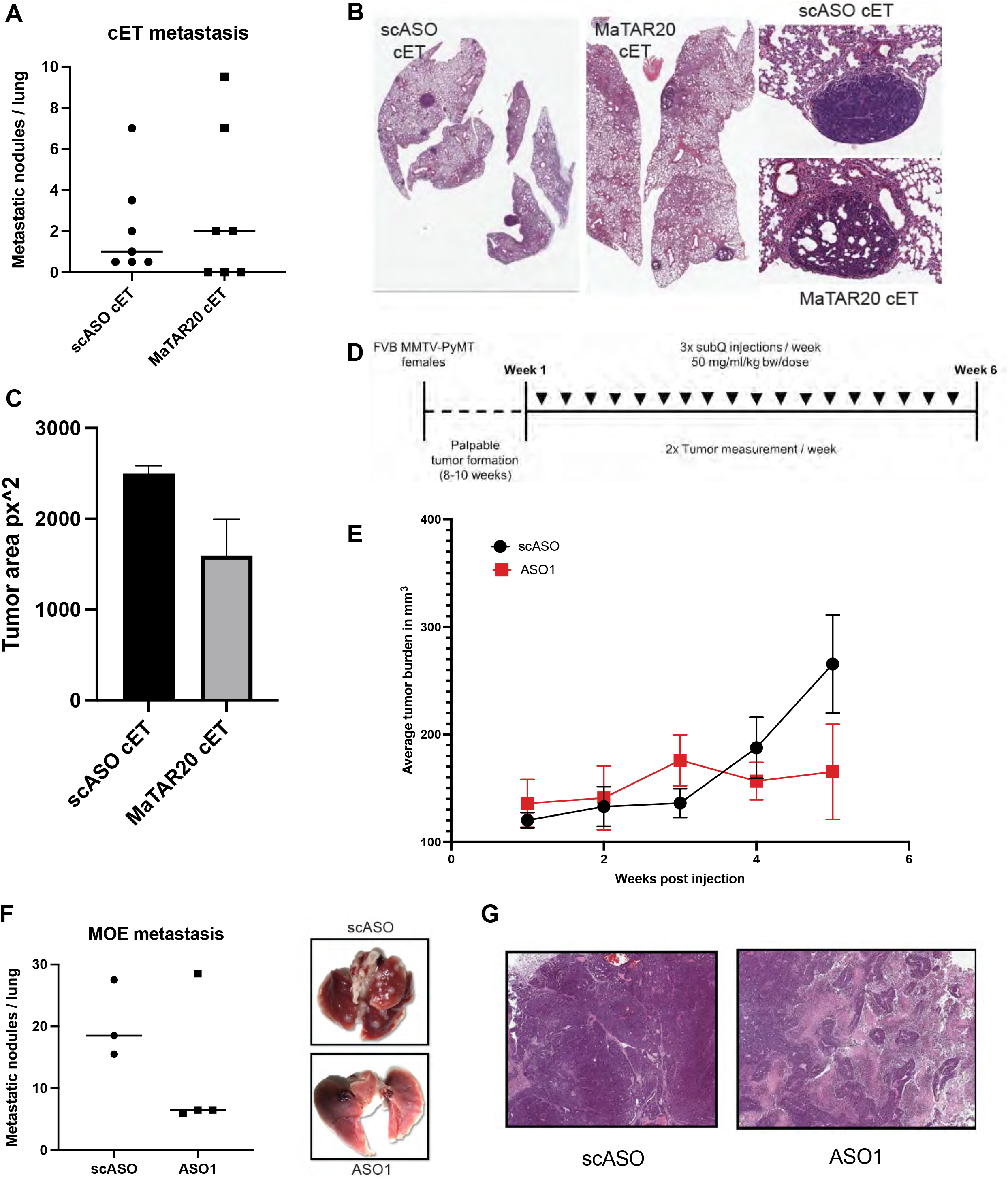
**A:** Related to Figure 5, cET experiments: quantification of macro-metastatic nodules, displayed is the average number per lung. **B:** Related to Figure 5, cET experiments: Hematoxylin and Eosin staining of fixed lung sections. **C:** Related to Figure 5, cET experiments: quantification of metastases by area. Bars represent the mean +/- SEM. **D:** MMTV-PyMT (FVB) mice were treated for 6 weeks with MOE ASOs (150 mg/kg/week), either scASO or *MaTAR20* ASO1. Bars / lines denote the mean of biological replicates +/- SEM. **E:** Average tumor burden in mm^3^ per week in each treatment group, tumor size 100-200 mm^3^, n=5 tumors. **F:** Quantification of macro-metastatic nodules, displayed is the average number per lung. Insets show exemplary images of lungs from scASO and ASO1 treated animals. **G:** Hematoxylin and Eosin staining of tumor sections. Left: scASO treated tumor. Right: *MaTAR20* ASO1 treated tumor.

**Figure S6:**
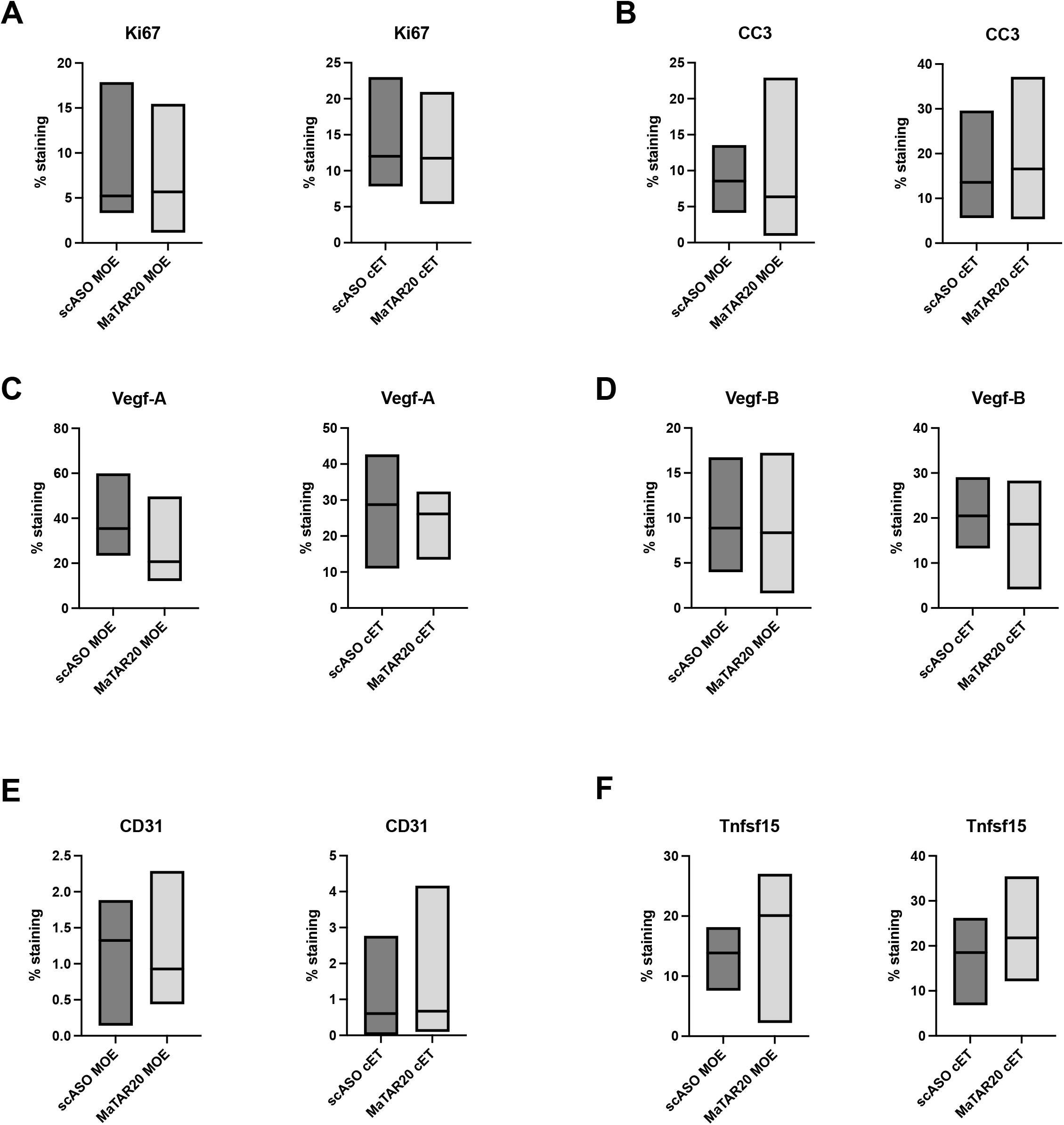
Immunohistochemistry on tumors from animals treated with either scASO or *MaTAR20* ASO. “MOE” = animals in the 2’MOE ASO cohort, “cET” = animals in the cET ASO cohort. Lines within boxplots indicate the median. The y-axis indicates the percentage of each tumor section stained with the respective antibody given a defined detection threshold (0.2 for CD31, 0.5 for all other antibodies). Two technical replicates were measured for each tumor. The axis indicates which ASO chemistry was used (MOE or cET). **A:** Ki67 staining **B:** Cleaved caspase-3 staining **C:** Vegf-A staining **D:** Vegf-B staining **E:** CD31 staining **F:** Tnfsf15 staining

**Figure S7:**
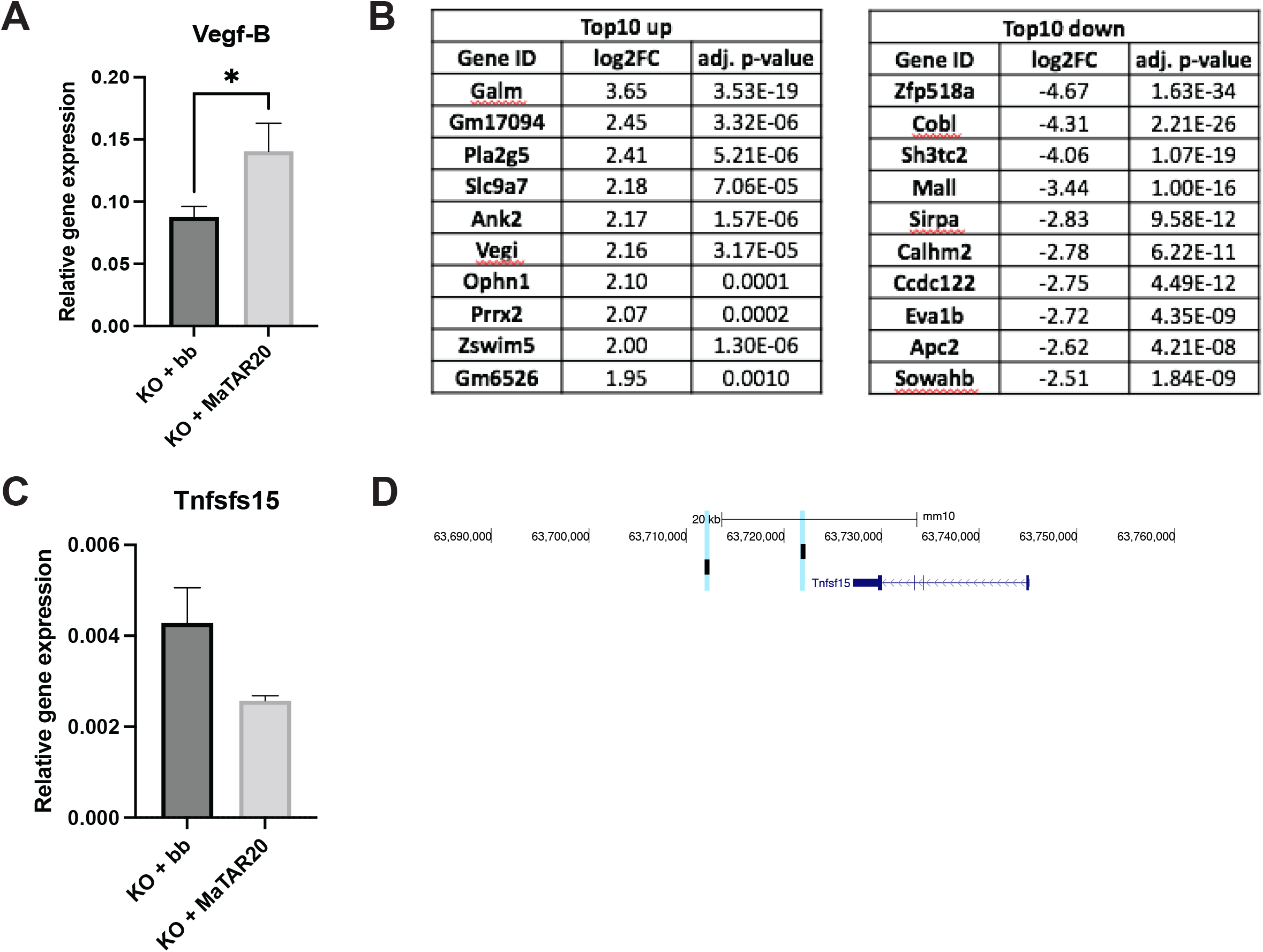
**A:** qRT-PCR to determine the relative *Vegf-B* expression in 4T1 *MaTAR20* KO cells. Cells were transfected either with *MaTAR20* Tx1 (KO+MaTAR20) or an empty backbone (KO+bb). Bars denote the mean of at least three biological replicates +/- SD. Statistical significance was determined with a two-tailed Student’s t-test; * p < 0.05 **A:** Top 10 up-and down-regulated genes in *MaTAR20* KO cells compared to Ren control cells. **B:** Like A, for *Tnfsf15*. **C:** UCSC Browser image illustrating *MaTAR20* binding to the *Tnfsf15* locus.

